# Human Phageprints: A high-resolution exploration of oral phages reveals globally-distributed phage families with individual-specific and temporally-stable community compositions

**DOI:** 10.1101/516864

**Authors:** Gita Mahmoudabadi, Kelsey Homyk, Adam Catching, Helen Foley, Arbel Tadmor, Ana Mahmoudabadi, Allison Cheung, Rob Phillips

## Abstract

Metagenomic studies have revolutionized the study of novel phages. However these studies trade the depth of coverage for breadth. In this study we show that the targeted sequencing of a phage genomic region as small as 200-300 base pairs, can provide sufficient sequence diversity to serve as an individual-specific barcode or “Phageprint”. The targeted approach reveals a high-resolution view of phage communities that is not available through metagenomic datasets. By creating instructional videos and collection kits, we enabled citizen scientists to gather ∼700 oral samples spanning ∼100 individuals residing in different parts of the world. In examining phage communities at 6 different oral sites, and by comparing phage communities of individuals living across the globe, we were able to study the effect of spatial separation, ranging from several millimeters to thousands of kilometers. We found that the spatial separation of just a few centimeters (the distance between two oral sites) can already result in highly distinct phage community compositions. For larger distances, spanning the phage communities of different individuals living in different parts of the world, we did not observe any correlation between spatial distance and phage community composition as individuals residing in the same city did not have any more similar phage communities than individuals living on different continents. Additionally, we found that neither genetics nor cohabitation seem to play a role in the relatedness of phage community compositions across individuals. Cohabitating siblings and even identical twins did not have phage community compositions that were any more similar than those of unrelated individuals. The primary factor contributing to phage community composition relatedness is direct contact between two habitats, as is demonstrated by the similarity between oral phage community compositions of partners. Furthermore, by exploring phage communities across the span of a month, and in some cases several years, we observed highly stable community compositions. These studies consistently point to the existence of remarkably diverse and personal phage families that are stable in time and apparently present in people around the world.

## Introduction

The study of bacteriophages, or viruses of bacteria, has traditionally relied on the culturing of the bacterial hosts. Because the vast majority of bacteria remain unculturable, we have only recently begun to recognize the overwhelming presence of phages through culture-independent techniques (1, 2). These advances collectively paint phages not only as the most numerous and diverse biological entities on our planet, but also as regulators of microbial ecosystems through rapid infection cycles and gene transfer events (3-7). Yet, compared to their bacterial hosts, and despite their proven potential to transform fields such as medicine (8-10), agriculture (11, 12), and biotechnology (13-15), phages are, in general, poorly characterized (16-20).

Even across familiar microbial habitats such as those within the human body, the identity of phages and their corresponding bacterial hosts, their community compositions, their modes of transfer between habitats, their co-evolutionary history with bacterial and human hosts, their role in health and disease, among other important topics remain highly unexplored. We thus chose to study the human oral cavity, not only because it represents a multifaceted and medically important ecosystem, but also because there are very few studies focused on oral phage communities (21-25).

Several intriguing studies have revealed phages as the most abundant members of the human oral cavity (10^8^ virus like particles per mL of saliva) (22, 26), with distinct communities at sites of disease (27, 28), capable of augmenting the bacterial arsenal of pathogenic genes (29, 30). These studies have relied on the shotgun metagenomic approach, in part because one of the defining features of viral genomes is the lack of a universally conserved sequence. Given that the ribosomal RNA sequence can be used as a universal marker for cellular genomes, its sequence variation is used to draw conclusions about cellular evolution and taxonomic classification (31-33). This marker-based approach to microbiology is additionally indispensible to microbial ecology as it allows a high coverage depth of the 16S region, which in turn, enables precise and reproducible depictions of bacterial community compositions (34-38).

Using the current sequencing platforms, the trade-off for coverage depth is typically the coverage breadth (Figure 1). In comparison to the marker-based approach, shotgun metagenomics provides a much greater breadth in coverage and offers several advantages. However, it suffers from several disadvantages. The coverage depth is often heterogeneous and remains comparatively low in these studies, a manifestation of which is that the *de novo* assembly of genomes from complex environments remains a significant challenge (39), even for abundant members with relatively short genome lengths (40). Moreover, the genomes assembled through shotgun metagenomics are often consensus genomes or an average representation of similar genomes within an environment (41). It is typical to see genomic segments with ∼100x coverage that are islands in the sea of lower coverage depth regions (42-44). Even across regions with high coverage depth, a notable limitation surfaces when there are variants that occur with a frequency below the detection limit.

**Figure 1.**
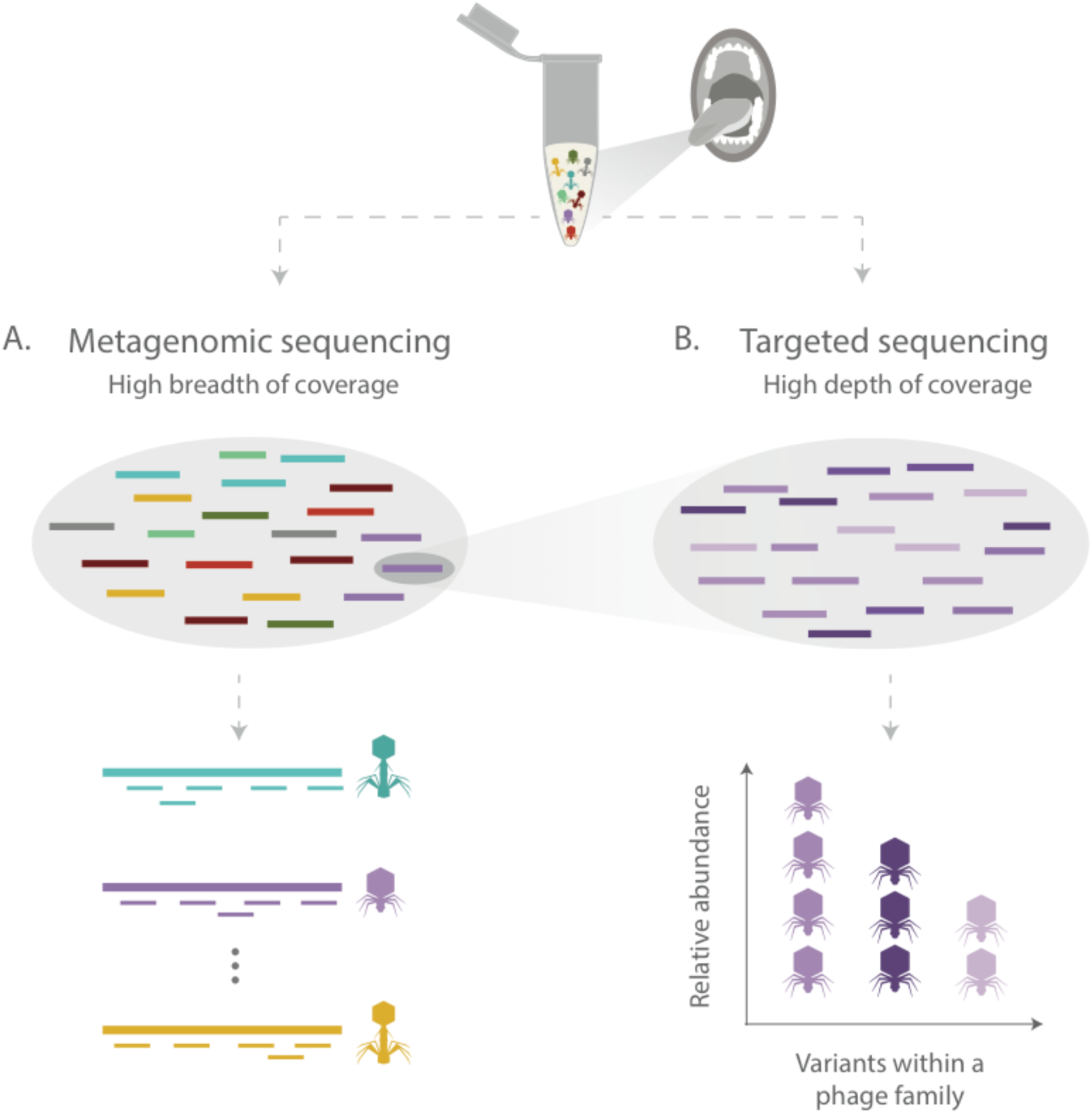
Comparison of A) shotgun metagenomic sequencing and B) targeted sequencing approaches. A) Shotgun metagenomic sequencing offers high breadth of coverage, spanning genomes from many different organisms, however it suffers from low depth of coverage (shown here by the incomplete assembly of phage genomes). B) Targeted sequencing approaches, which use PCR to amplify a specific genomic region, exchange breadth of coverage for depth. Targeted sequencing studies, due to their greater depth of coverage, provide much higher resolution for constructing the community composition by equating coverage depth with relative abundance of species or strains.

Due to these technical challenges, the marker-based approach, which allows orders of magnitude greater coverage depth by focusing the reads on a small genomic segment, provides a higher resolution view of community compositions. The targeted approach is therefore widely used to complement shotgun metagenomic depictions of bacterial communities (45-47).

Due to the immense sequence diversity of phage genomes (48-50), an in-depth view of their communities through a targeted sequencing approach could provide novel insights. As such, the overarching aim of this study was to explore oral phage communities and their inter-and intra-personal diversity, their spatial patterns of distribution, as well as temporal dynamics in a large-scale and high-resolution fashion. Towards this aim, we first had to choose regions within phage genomes on which to perform targeted sequencing. Because of the vast diversity of uncultured phages, we relied on oral metagenomic datasets to identify candidate marker sequences from such phages.

We have described the methods for phage marker discovery and validation in our recent manuscript (Tadmor *et al*., in preparation). Briefly, we arrived at seven phage markers, each corresponding to a distinct lineage of the terminase large subunit gene (TerL), which is involved in the packaging of DNA inside the phage capsid. The terminase was chosen as a target because it is considered to be a uniquely phage gene encoded by many double-stranded DNA phages (51, 52). In the absence of a genomic taxonomy for viruses, we will refer to those phages (or prophages) that share similar TerL sequences as members of a phage family. This assumption is predicated on previous studies that have shown no significant sequence similarity between TerL sequences of unrelated phages (53-55). In this study we will explore three of the seven markers, and will refer to those phages that contain these TerL markers as members of the HA, HB1 or PCA2 phage families. Refer to Materials and Methods for further information on marker discovery.

By designing primers to target these phage families, we were able to obtain at least several thousand sequences per marker, per individual (see Materials and Methods). As a direct comparison to our exploration of the same markers from hundreds of shutgun metagenomic samples (Tadmor *et al*., in preparation), this study increases the coverage depth of a marker per subject by at least three orders of magnitude (from a few sequences to a few thousand sequences). We will demonstrate that at high sequencing depth, the phage community composition derived from members of just a single phage family can already serve as a fingerprint, or a “phageprint” – highly unique to a microbial habitat and stable over time.

By creating instructional videos and collection kits, we enabled citizen scientists to gather ∼700 oral samples spanning ∼100 individuals residing in different parts of the world. As a point of comparison, one of the largest studies of the human microbiome recently reported on data from 265 individuals (56). By examining phage communities at 6 different oral sites, and by comparing phage communities of individuals living across the globe, we were able to study the effect of spatial separation, ranging from several millimeters to thousands of kilometers. We found that the spatial separation of just a few centimeters (the distance between an individual’s gingival sites and the hard palate, for example) can already result in highly distinct phage community compositions. For larger distances, spanning the phage communities of different individuals, we did not observe any correlation between spatial distance and phage community composition. In other words, individuals residing in the same city did not have any more similar phage communities than individuals living on different continents.

Additionally, we found that neither genetics nor cohabitation seem to play a role in the relatedness of phage community compositions across individuals. Cohabitating siblings and even identical twins did not have phage communities that were any more similar than those of unrelated individuals. The only factor we observed that contributes to phage community relatedness is direct contact between two habitats, as is demonstrated by the similarity between oral phage community compositions of partners. Furthermore, by exploring phage communities across the span of a month, and in some cases several years, we observed highly stable communities. These studies consistently point to the existence of remarkably diverse and personal phage families that are stable in time and apparently present in individuals living in different parts of the world.

## Methods Summary

From a methodological standpoint, targeted sequencing of these phage markers is very similar to 16S sequencing (35, 57). Using barcoded primers, we employ PCR and next generation sequencing to attain millions of paired-end reads (Figure 2). After several quality control filters, the reads are demultiplexed based on their barcoded primer sequence and linked back to the sample and the marker from which they originated. All reads derived from the same primer sets (i.e. reads corresponding to the same TerL marker) are then pooled and clustered based on their sequence similarity into Operational Taxonomic Units or OTUs. An OTU table is constructed wherein the number of reads belonging to each OTU across each sample is denoted. Using the OTU table, we can plot the relative abundance of each OTU within a sample. We refer to this plot, which represents relative abundance profile of members within a phage family as a “community composition” plot. Finally, we will use various diversity metrics to further explore phage communities within and between individuals. This procedure is repeated for each of the three phage families. Detailed description of these protocols can be found in the Materials and Methods.

**Figure 2.**
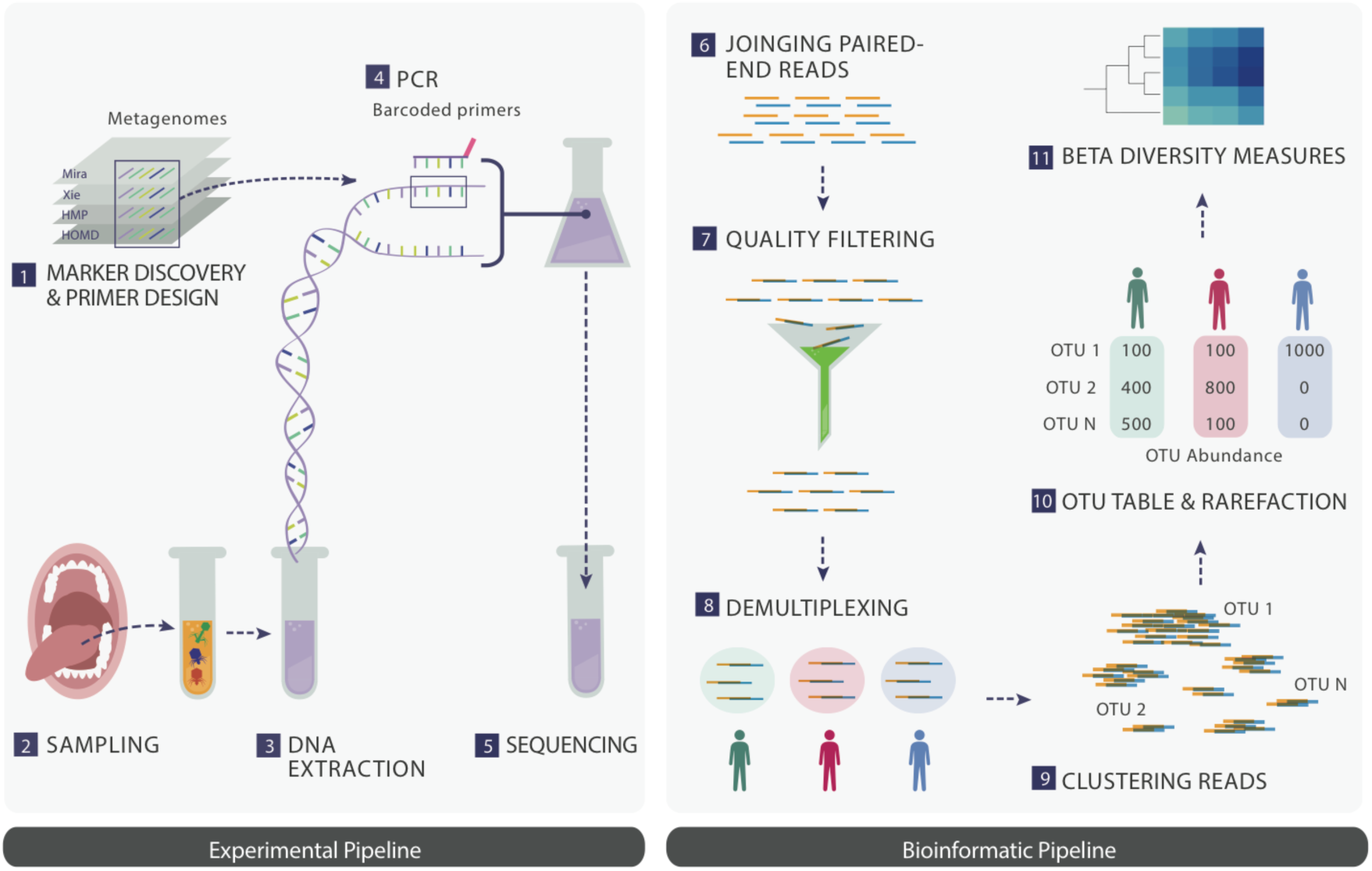
A schematic summary of the main experimental and bioinformatic methods: 1) Discovery of ubiquitous phage families by examining large terminase sequences that occur across different metagenomic datasets, 2) experimental sampling, 3) DNA extraction from oral biofilm samples, 4) PCR using barcoded primers followed by PCR clean-up, 5) paired-end Next Generation Sequencing, 6) joining paired-end reads to eliminate sequencing errors, 7) additional quality control steps to further eliminate errors, 8) demultiplexing of reads based on their barcode sequence and linking sequences to the sample they originate from, 9) gathering reads from all samples and clustering them based on sequence similarity (OTUs), 10) counting the number of sequences belonging to each OTU from each sample (i.e. constructing an OTU table), and rarefying the table so that each sample is represented by the same total number of sequences, and 11) performing various downstream diversity analysis (e.g. community composition plots or phageprints) using the constructed OTU table as the basis. Note that these steps are performed separately for each of the three phage families (three separate OTU tables are constructed).

## Results

### An exploration of phage families reveals the presence of highly diverse, personal phage community compositions that are stable in time

As previously described in the introduction, due to the high depth of coverage afforded through targeted sequencing, we are able to explore phage sequence diversity in extraordinary detail. With bacterial 16S data, sequences are generally clustered at 97% sequence similarity into operational taxonomic units (OTUs), primarily to manage the large volume of data. At this threshold, each OTU is conventionally referred to as a bacterial species. In fact, it is based on OTU counts that the number of bacterial species in a habitat is estimated. In the absence of any convention for handling viral targeted sequencing data, we have used here various sequence similarity thresholds for clustering (including 100% sequence similarity threshold). We found the results to be largely robust to variations in the sequence similarity threshold (see Materials and Methods).

Figure 3 demonstrates the HA phage community composition from a subject’s tongue dorsum (top surface) at two time points. The x-axis is a list of OTUs that were generated when HA phage family sequences from all subjects were clustered based on sequence similarity. The y-axis counts the relative abundance of the subject’s HA sequences that fall into each OTU. As shown in this representative figure, and across all other community composition plots we have seen for all three phage families, the community composition is highly skewed towards a small number of dominant OTUs (typically one or two OTUs). In addition to these OTUs, there are many other OTUs with abundance values that are fairly stable in time. Generally, the dominant OTUs are not the same across different individuals, and the presence of numerous other OTUs with temporally stable relative abundances, gives rise to phage community compositions that are highly personal. Therefore, we coined “phageprint” as shorthand to refer to a community composition plot.

**Figure 3.**
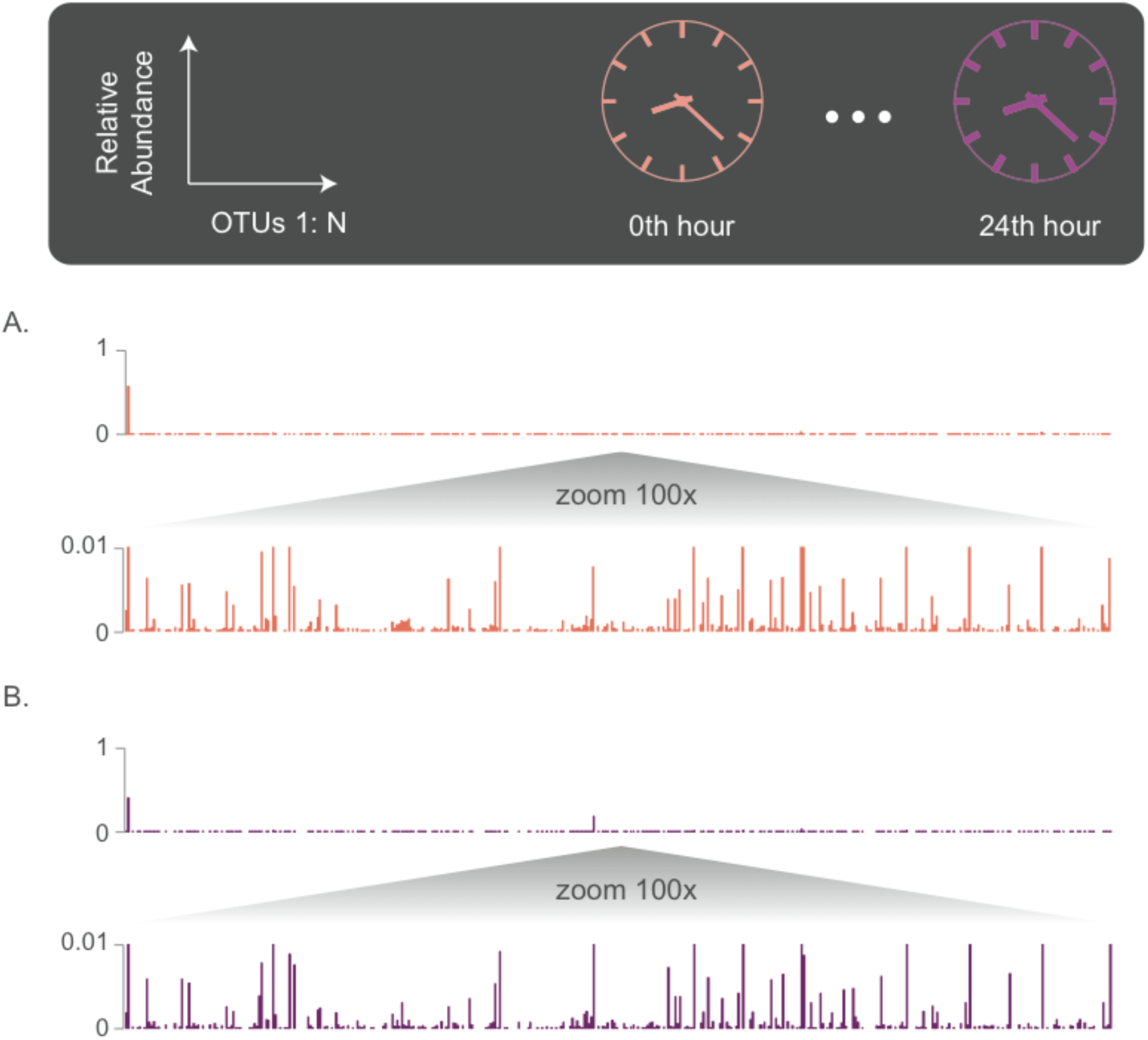
HA Phage community compositions (phageprints) from subject 37 at two different time points. Samples were collected from the tongue dorsum. A) Subject 37’s phageprint at 0^th^ time point, collected right after brushing tongue dorsal and teeth surfaces. B) Subject 37’s phageprint 24 hours after the initial time point (no brushing in between time points). Each phageprint is derived from the analysis of 4000 sequences. OTUs are defined at 98% sequence similarity.

We have so far demonstrated the highly personal nature of phage communities residing in the human mouth. To better explore the temporal dynamics of these phage communities, 10 subjects collected biofilm from the tongue dorsum every 24 hours for 30 days. The HB1 community composition as it evolved over 30 days on subject 1’s tongue dorsum is depicted in Figure 4. Here, to provide a more detailed view of this community, we cluster the HB1 TerL sequences into OTUs based on 100% similarity.

**Figure 4.**
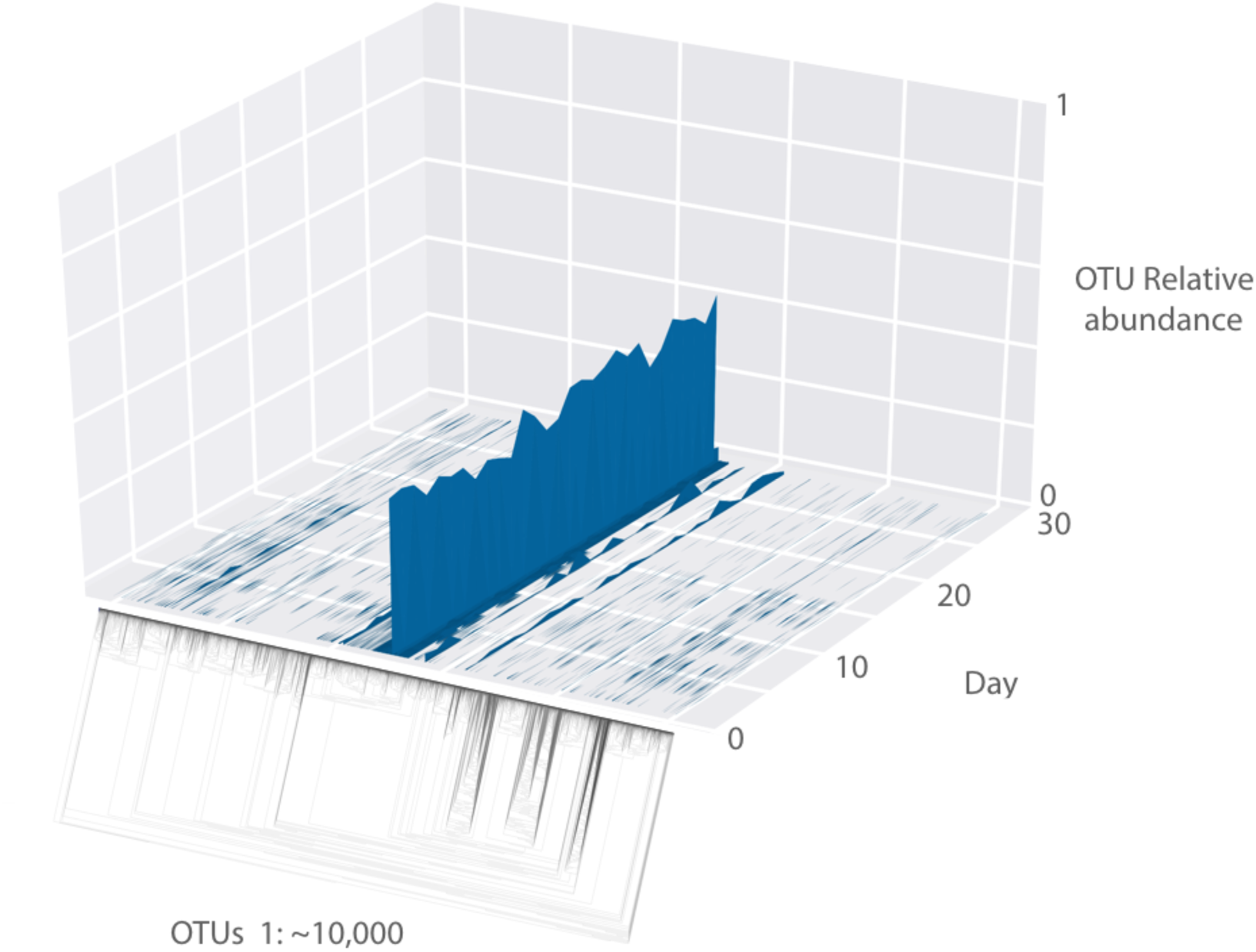
A 3D surface plot depicting the HB1 phage community composition as it evolves over 30 days on subject 1’s tongue dorsum. The x-axis contains ∼10,000 OTUs ordered according to the depicted phylogenetic tree of the OTU sequences (the phylogenetic tree is provided largely to serve as a schematic since it is hard to visualize the details of this tree). Each OTU is composed of identical sequences (i.e. 100% sequence similarity threshold). The y-axis depicts the relative abundance of each OTU, and the z-axis shows the fluctuations in relative abundance of each OTU in time.

Surprisingly, over 30 days, the main features of each phage community composition is preserved, though there are also interesting fluctuations that are well above the experimental error and detection threshold (see SI). Figure 5 demonstrates different degrees of temporal stability and phylogenetic diversity across individuals. However, a global trend is that the dominant OTU(s) remain dominant over the span of 30 days in all subjects. This observation is especially interesting in light of the inter-and intra-personal differences in diet and oral hygiene practices over time (Figure 6).

**Figure 5.**
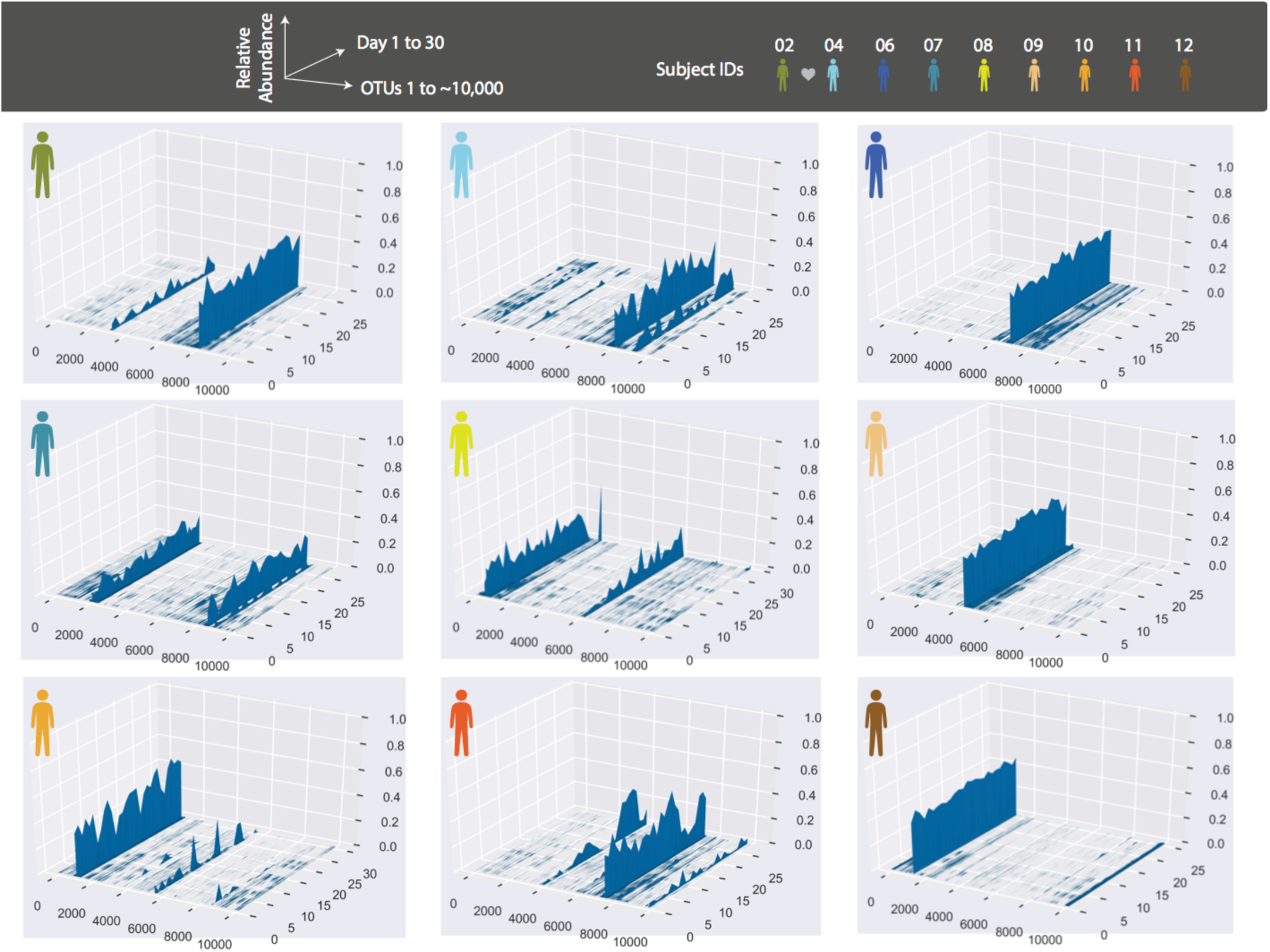
Depictions of HB1 phage community composition evolution in different subjects over 30 days. The format of the plots is the same as that of Figure 4, and the order of OTUs is based on their phylogenetic distance and identical across all plots. All samples are collected from the tongue dorsum. Note that subject 2 and 4 are a couple, and their phage community compositions share some main features. The metadata associated with these subjects is provided in Figure 6.

**Figure 6.**
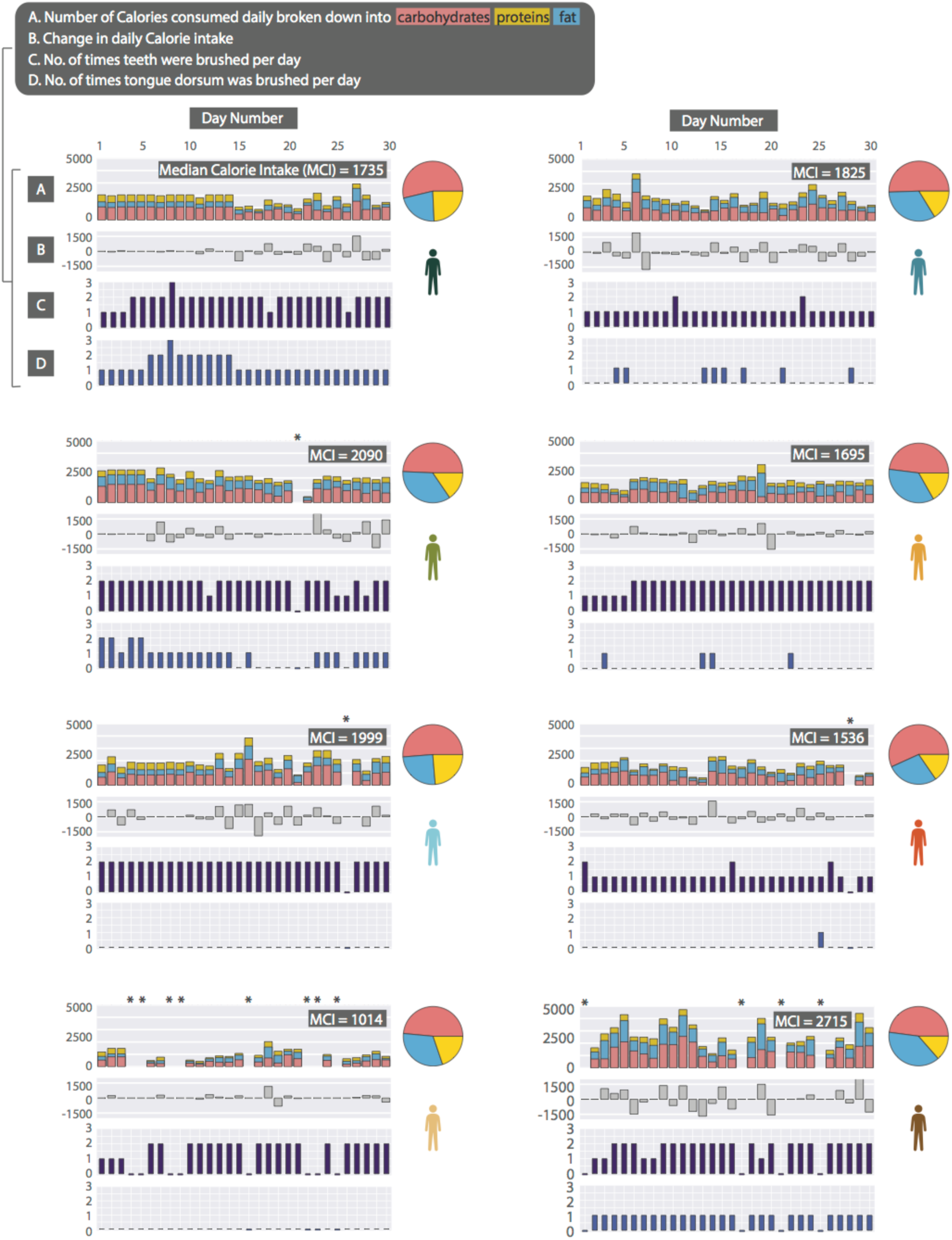
Subject daily metadata during 30 consecutive days. Top panel for each subject represents the Caloric intake from fats, carbohydrates, and protein. Mean Caloric intake or MCI reports the Caloric intake averaged over 30 days (the x-axis for all plots is number of days). Pie charts demonstrate the diet over 30 days based on median fat, carbohydrate and protein consumption. The second panel depicts the change in Calorie intake from the previous day. The third and fourth panel correspond to the number of times that the subject brushed his or her teeth and tongue, respectively, during the 24 hour sampling interval. We have used asterisk to denote days for which we did not receive data from subject, and to distinguish them from zero values in third and fourth panel, they have been given “-.1” value. The subjects are the same as those shown in the previous two figures, however two of the subjects did not report dietary information so they are not included in this figure.

To make quantitative pairwise comparisons between community compositions we employed several commonly used metrics such as the Bray-Curtis and Unifrac (see Materials and Methods), and in doing so, we distill the comparison of thousands of sequences from any two samples to a single score. We will therefore present heatmaps of pairwise comparison scores for each phage family.

All distance metrics explored paint similar pictures of the HB1 phage communities, depicting them as highly personal and stable over time (Figure 7, Figure 8). Because phage communities in different individuals have such distinct compositions, abundance-based metrics are especially suitable for describing them. However, even the binary Jaccard and weighted Unifrac distance metrics demonstrate a similar message. Figure 8 further demonstrates the intra-and -interpersonal distances as measured through these various distance metrics. As is expected from the heatmaps shown in Figure 7, the intra-personal distances are markedly different from the inter-personal, with the notable exception being subject 2 and 4, who are partners.

**Figure 7.**
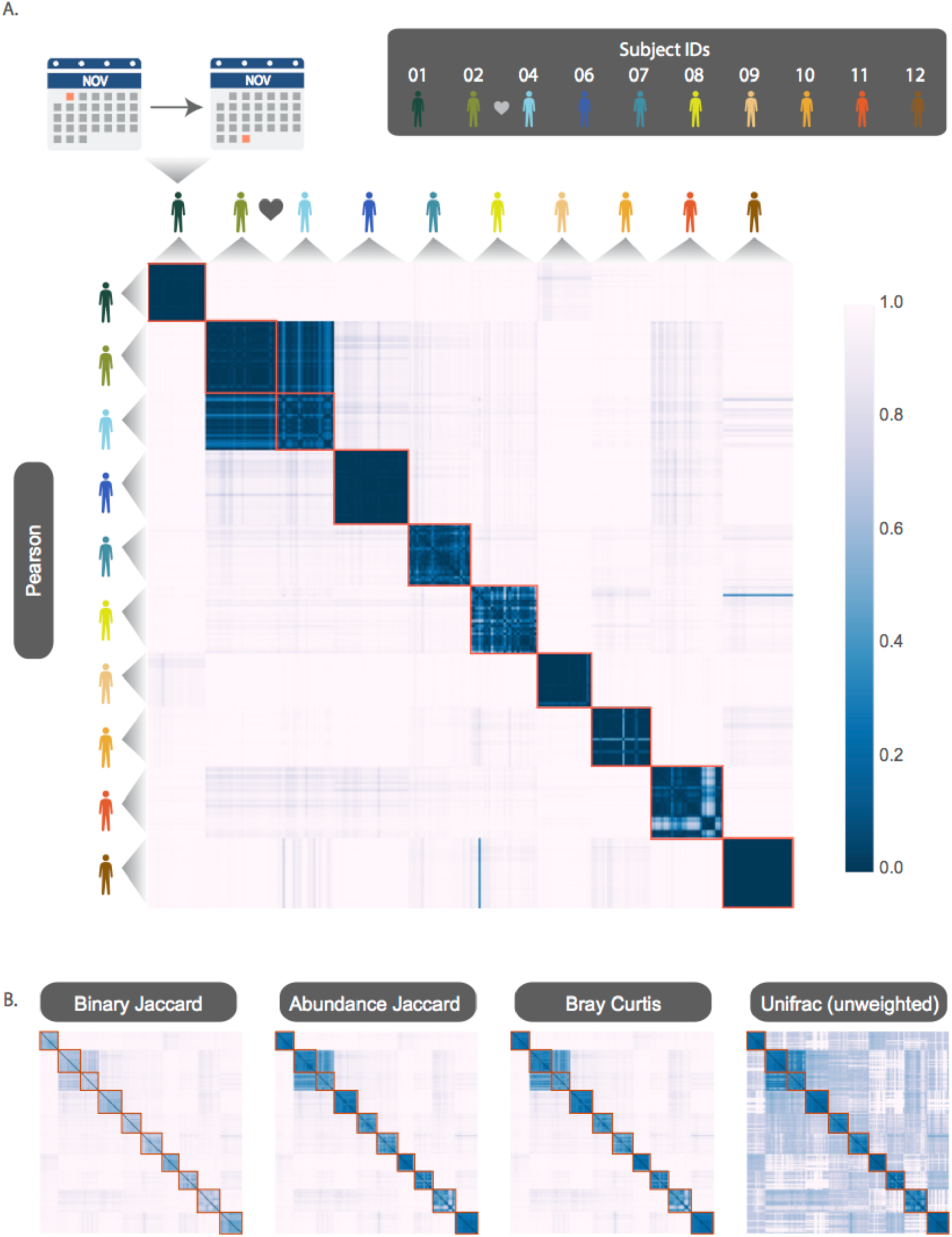
HB1 phage community temporal dynamics (previously shown graphically in Figure 5) depicted here by pairwise distance metrics: A) Peason, B) Binary Jaccard, Abundance Jaccard, Bray Curtis and unweighted Unifrac. The heatmap scale applies to all heatmaps shown. Subjects 02 and 04 are a couple. Samples from each subject are chronologically ordered.

**Figure 8.**
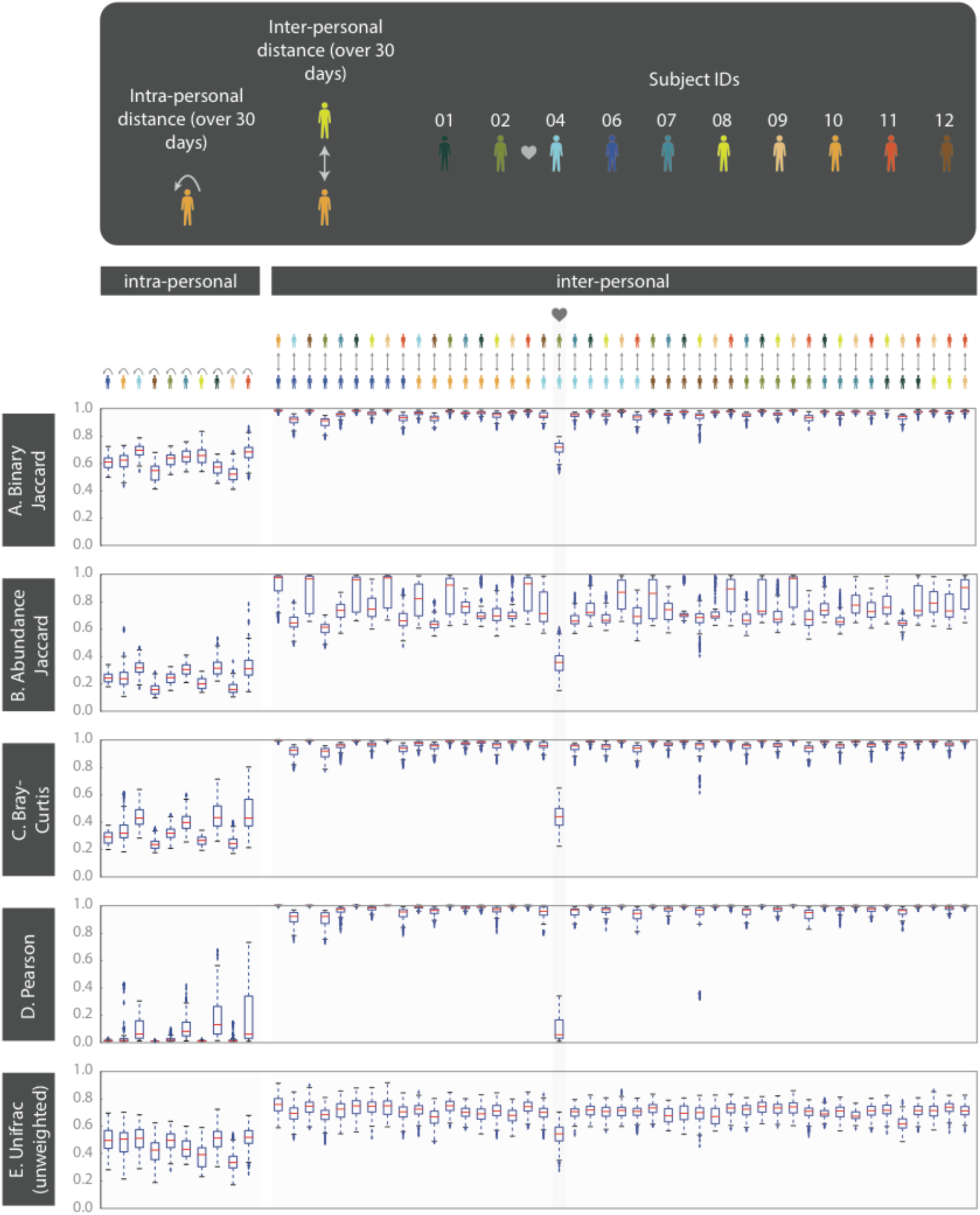
Intra-and inter-personal distances between HB1 phage communities from 10 subjects, over the span of 30 days (further quantifying the heatmaps from Figure 7). Box-plots depict distances from pairwise comparisons made using the following metrics: A) Binary Jaccard, B) Abundance Jaccard, C) Bray-Curtis, D) Pearson, and E) unweighted Unifrac. The outliers defined as those outside of the 1.5 x IQR (inter-quartile range) are denoted by “+”. The box-plots corresponding to the comparisons between the couple in this study are highlighted.

### Phage community comparisons across siblings, couples, and non-related individuals residing across the globe

Given the ubiquitous presence of the phage families across subjects residing in the U.S. we wondered whether phage families (HA and HB1, specifically) are globally distributed, and whether subjects residing in the same country would have more similar phage communities. We discovered that phage families were in fact found in individuals from various ethnicities, nationalities, and ages. Surprisingly, neither from abundance-based nor phylogenetic distance comparisons did we find an indication that people residing in the same country share more similar phage communities (Figure 9). Instead, we continued to find that individuals typically have highly unique phage communities.

**Figure 9.**
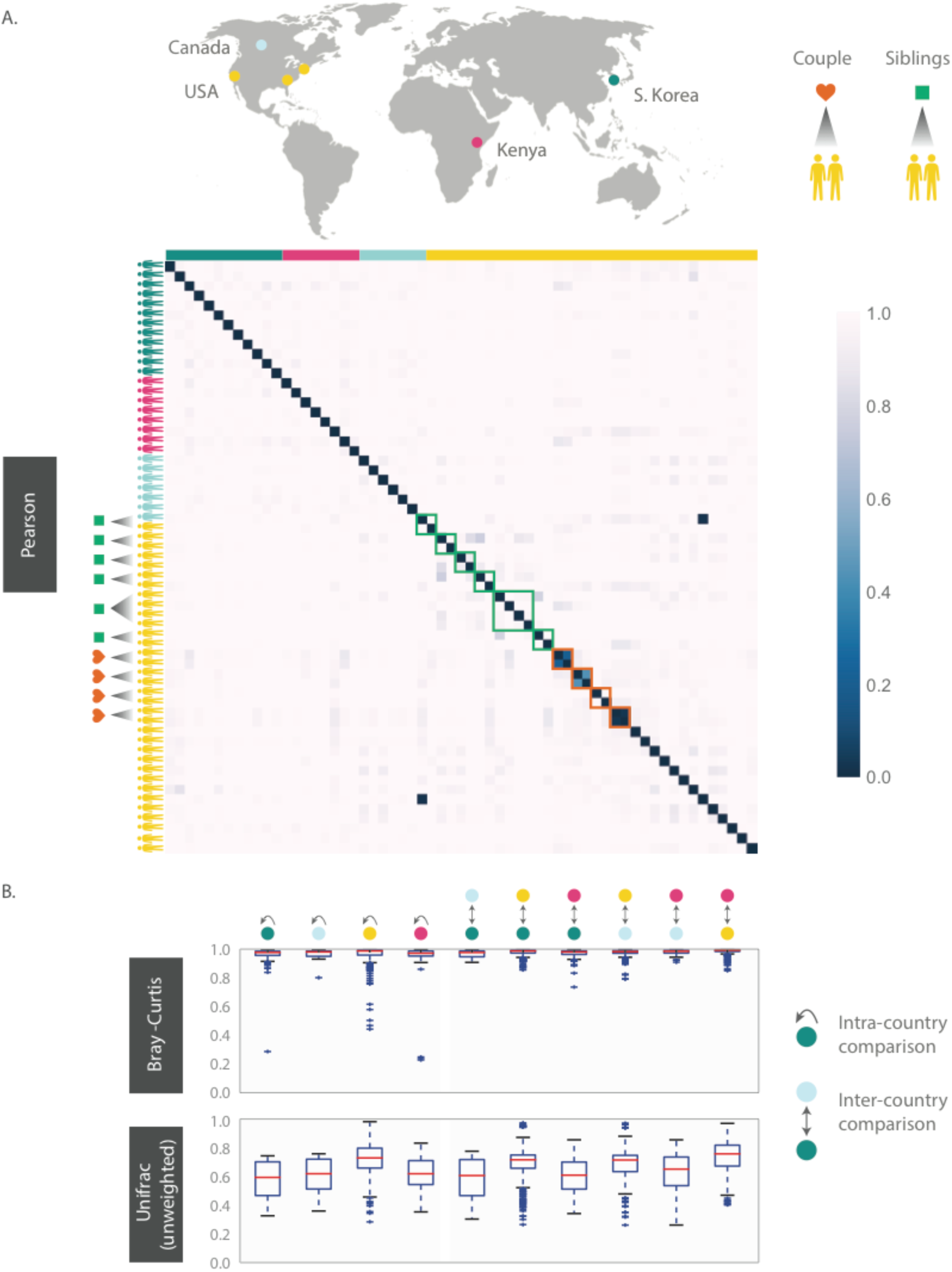
HB1 phage community across 61 individuals residing across different parts of the globe. Samples are obtained from the tongue dorsum. A) Pearson distance (1– Pearson correlation) is shown as a heatmap. A subset of individuals residing in the U.S. are either couples or siblings. Green and red boxes are drawn around samples from each sibling group and couple, respectively. B) Intra-and inter-country distances from pairwise comparisons made using Bray-Curtis and unweighted Unifrac distance metrics. The outliers are denoted as points outside of the 1.5 x IQR (inter-quartile range). Siblings and couples are excluded from this analysis.

Even siblings who were either living in the same household or had previously, do not have any more similar phage communities than unrelated individuals. In fact, one of the four sibling groups with uncorrelated phageprints are identical twins (Figure 9). However, 3 out of 4 couples in this study exhibited highly similar phage communities. The dissimilar couple may be due to celiac disease diagnosed in one of the partners, which is known to alter oral ecology (58). These results suggest that genetics and cohabitation do not significantly impact a person’s oral phage community. The more impactful factor appears to be direct oral contact with another person. To further test these trends, larger studies encompassing a greater number of individuals and regions in the world are required.

### Different oral sites

Thus far, all phage communities shown are those sampled from the tongue dorsum. In order to examine the spatial patterns of phage communities we obtained additional oral samples spanning 9 individuals and 6 oral sites (Courtesy of Bik *et al*.). Figure 10 shows the HB1 phage community compositions of a subject at four oral sites where HB1 phage family was found. Clearly, different oral sites in this subject have very similar HB1 phageprints. When examining all HB1 positive samples, an immediately recognizable pattern is that the HB1 phage community compositions of an individual are highly correlated. In stark contrast are the correlations between the phage community compositions of different individuals.

**Figure 10.**
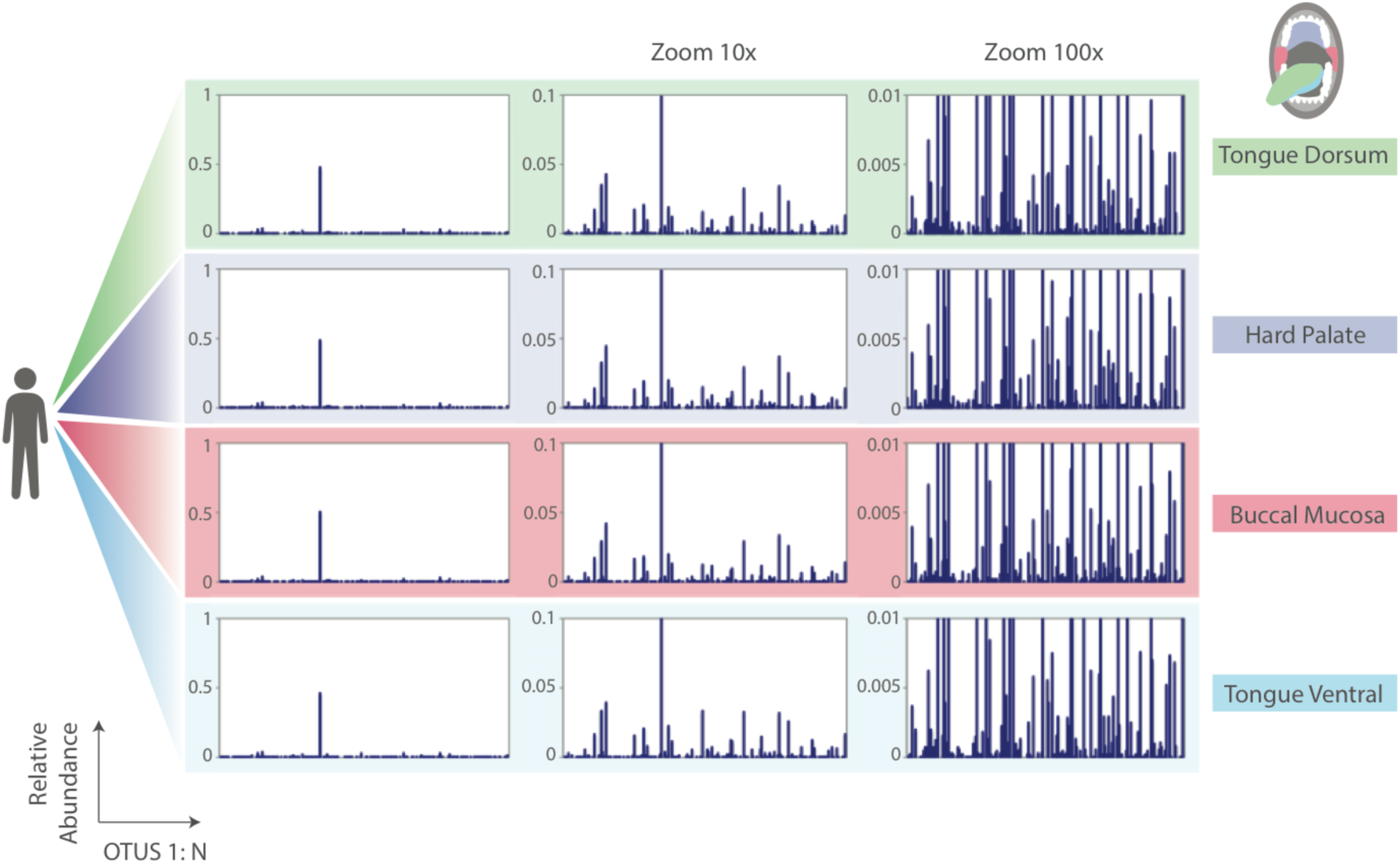
HB1 phage community compositions (phageprints) across 4 different oral sites in subject 16. Each phageprint is derived from the analysis of 4000 sequences (see SI). OTUs are defined at 98% sequence similarity and OTUs with less than or equal to 0.1% relative abundance across all phageprints were filtered out (see SI).

As in the case of the HB1 phage family, there is low to non-existing correlation between the HA phage community compositions of different individuals at the same oral site (Figure 12), reinforcing the notion of highly personal phage communities. However, unlike HB1, not all oral sites within the same subject are highly or even moderately correlated (see subjects 3, 12, and 17). In subject 12 for example, the tongue dorsum has a correlation close to zero with supra-gingiva and sub-gingiva sites, which are nearly perfectly correlated. Similarly, in subject 3, the hard palate and the tongue ventral surface have nearly identical phage community compositions while they have a very low correlation with the community at the tongue dorsum. However, unlike subject 12, the tongue dorsum in subject 3 seems to be an intermediate community, having a moderate correlation with all other sites that are distinct from each other. In subject 17 as well, buccal mucosa serves as the intermediate community, having a moderate correlation with the disparate communities of sub-gingiva and the hard palate. Phage-host network representations for HB1 (SI Figure 1) and HA (SI Figure 2) phage families across this cohort demonstrates in extensive detail the cause of weak or strong correlations between different oral sites.

**Figure 11.**
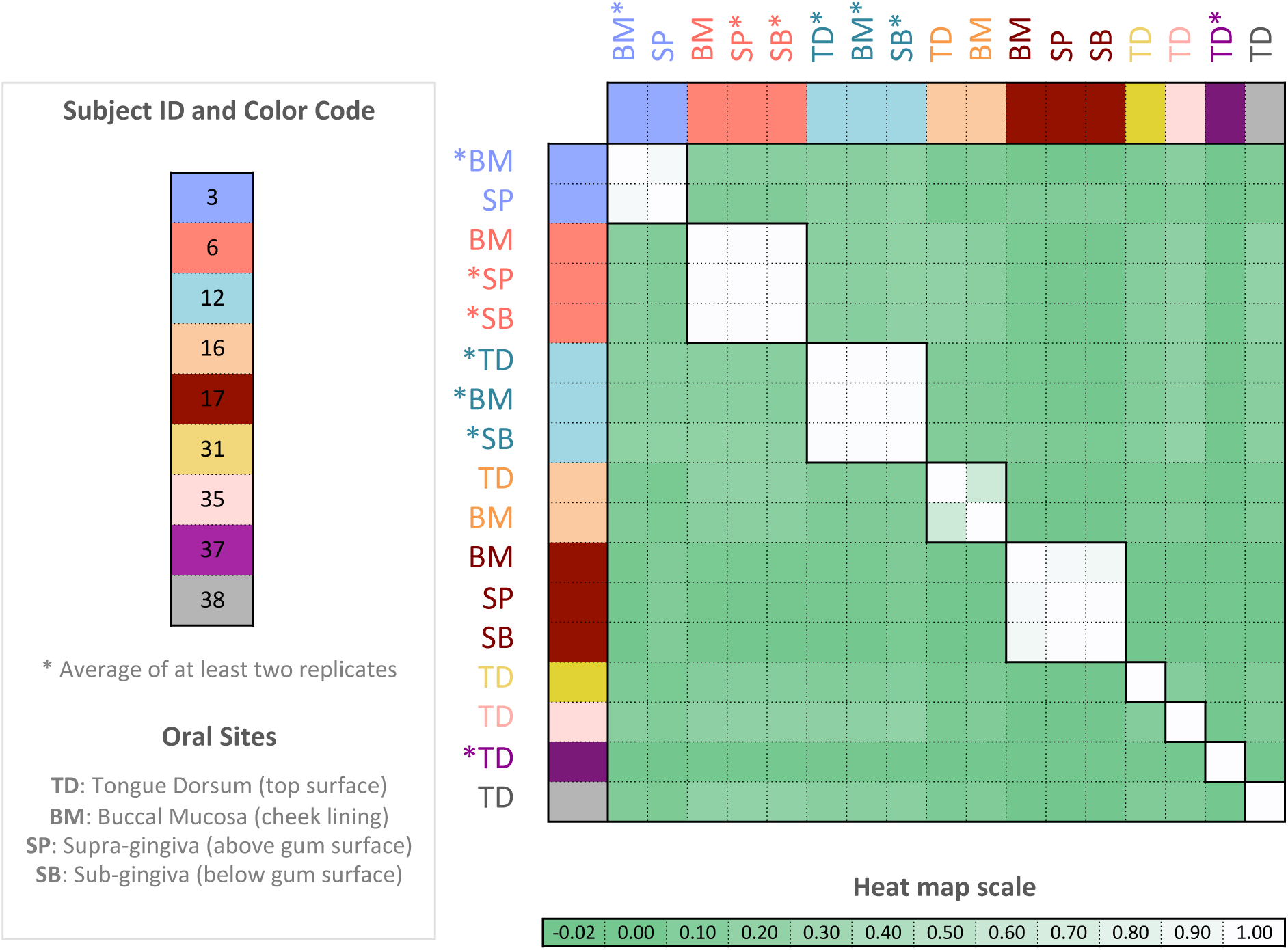
Pearson correlation coefficient matrix of HB1 phage community compositions spanning 9 subjects and four oral sites. Each community composition is derived from the analysis of 4000 sequences associated with an individual and a particular oral site. OTUs are defined at 98% sequence similarity and OTUs with less than or equal to 0.1% relative abundance across all phageprints were filtered out (see SI). Phageprints are color-coded based on the individual they originate from. Community compositions that have been replicated at least twice and averaged have an asterisk next to them (see SI).

**Figure 12.**
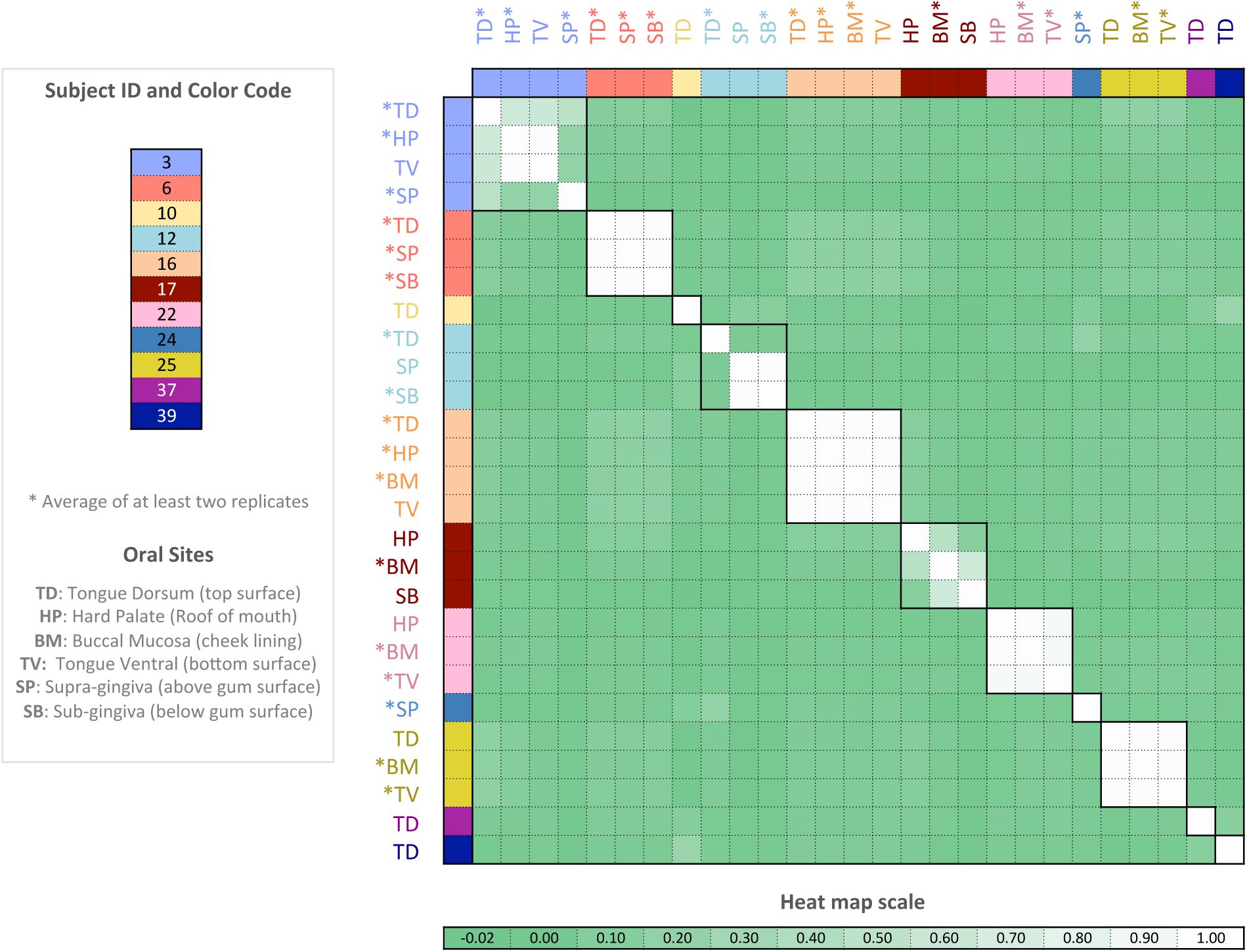
Pearson correlation coefficient matrix of HA community compositions encompassing 11 subjects and six oral sites. Each community composition is derived from the analysis of 4000 sequences associated with an individual and a particular oral site. Samples are color-coded based on the individual they originate from. Oral sites shown are the tongue dorsum (TD), buccal mucosa (BM), supra-gingiva (SP), sub-gingiva (SB), hard palate (HP), and ventral surface of the tongue (TV). Samples whose community composition has been replicated at least twice and averaged have an asterisk next to them (see SI).

### A bioinformatic search for the bacterial hosts

Because we aimed to study previously uncharacterized phages, the bacterial hosts for the phage families in this study have not yet been cultured or identified. However, using homology-based searches we can identify candidate host species. For each phage family, the most abundant sequence in each OTU served as its representative sequence and was used as a query for BLASTx homology search. With the exception of a few sequences tagged as “putative proteins”, all resulting homologs were terminase sequences (SI Table 1, SI Table 3, SI Table 5). Additionally, the bacterial species with the highest chance of harboring members of these three phage families were determined based on the results of the BLASTx homology search (SI Table 2, SI Table 4, SI Table 6, SI Figure 3).

The HA phage family infects only a single genus of Firmicutes (*Streptococcus*), but appears in the genomes of many different species within this genus (SI Table 4). The majority of HB1 homologs belonged to ReqiPepy6 phage isolated from *Rhodococcus equi* (phylum Actinobacteria). Other OTU homologs were matched to ReqiPoco6, another *R. equi* phage, and six species spread across two different families of the Firmicutes phylum.

## Discussion

Our method for finding ubiquitous human oral phages relied on a relatively small metagenomic dataset, which contained sequences from 6 individuals residing in Spain (59). Yet, on the basis of markers designed from this small dataset we were able to identify the same phage families in at least 10 times as many individuals from across the globe. This finding seems to suggest that there exist certain phage families that are a stable feature of the human oral microbiome. Studies of phages from various natural environments (e.g. marine, soil, lakes) also report the finding of phage families that are distributed across similar types of habitats despite vast geographical distances and barriers that exist between these habitats (17, 60, 61). The discovery of core bacterial members within the human microbiome (21, 62, 63) that are present globally (23) further support our discovery of globally distributed phage families. Similar to our findings for phages, the oral bacteria of individuals from the same part of the world was as different from each other as they were to individuals from other parts of the world (23).

The ubiquitous presence of the identified phage families in individuals, together with their temporal stability, seems to suggest that they likely play important roles in this environment. The observed temporal stability of these phage community compositions over the span of a month is supported through metagenomic studies of oral phages (64, 65) as well as 16S sequencing of bacterial communities inhabiting various sites in the human body (62, 66, 67). Our study represents one of the largest studies of human oral phages. As a comparison, the most recent version of the Human Microbiome Project, contains samples from 265 individuals (56). However, future studies are required to expand our dataset to include many more individuals and many more parts of the world.

A particularly important aspect of our study is that it combined the advantages of metagenomics with targeted sequencing to not only identify core phage families inhabiting the human oral cavity, but to also characterize their communities with a resolution that is unavailable through metagenomic studies of phages. This detailed view allowed us to clearly observe the highly complex, and personal nature of phage community compositions. Moreover, the emergence of phageprints is directly the result of the remarkable phage sequence diversity that we were able to capture via targeted sequencing. For example, we observed a few hundred HB1 OTUs (defined at 97% sequence similarity). Even though the HB1 phage family is only one of many oral phage families, it contains the same level of sequence diversity as the entire bacterial population in the human mouth. This perhaps explains the need to use highly elaborate algorithms applied onto 16S and metagenomic sequences from all bacterial strains to be able to identify a person based on his/her microbiome (68) whereas the personal nature of phage community composition plots is a clearly apparent feature of the datasets. Although it is widely known from shotgun metagenomic studies that viruses are highly diverse, targeted studies employing different types of bacterial and viral markers could perhaps enable more quantitative comparisons of sequence diversity across these organisms.

Because of the great diversity of sequences associated with just one phage family, in our study of about 60 individuals we found one case where unrelated individuals whose phageprints had similar correlation coefficients. This may be due to experimental error or due to the course-graining associated with Pearson correlation matrices. However, if we conservatively assume that each phage family can provide just 50 unique patterns, then the combination of phageprints from just 6 independent phage families would already provide a greater number of possible patterns than the size of the current human population. Identifying individuals based on characteristic patterns of anonymized personal microbiomes is a legitimate ethical concern (69) and should be considered with the potential uses of phageprints as well. Future studies are needed to test the long-term stability of the human phageprints especially with regard to perturbations such as exposure to antibiotics. To our knowledge, this is the first study that demonstrates the potential application of phage sequences for human identification.

## Materials and Methods

### Phage marker discovery

In our search for candidate phage markers, we combined the advantages of metagenomic and marker-based approaches. To study previously unexplored phage families, we first used existing metagenomic datasets to identify candidate phage families, and then by targeting these families using PCR, we were able to explore them with high depth of coverage. We limited our bioinformatic and later experimental search for ubiquitous phage families to those inhabiting the human oral cavity.

We have described our method for finding candidate phage markers (Tadmor *et al*., in preparation), but here we will provide a brief summary. Depending on the question of interest, there are potentially numerous ways by which phage marker sequences can be selected. In our method, we imposed several search criteria: 1) candidate markers should be unique to phages; 2) candidate markers should not share any significant sequence similarity so that they are more likely to represent distinct phage groups, and 3) candidate markers should be present across different oral metagenomic datasets so that they are more likely to represent core (though not necessarily abundant) members of the human oral phageome. Just as one gut phage genome was detected and assembled by an analysis that took into account its presence across multiple metagenomic datasets (40), we suspected that presence of a phage marker across multiple metagenomes could suggest that it belongs to a ubiquitous phage family. In the absence of a taxonomic convention for viral genomic data, we use the term “phage family” to refer to phages that share a given marker.

To meet the first criterion, we focused our search on the terminase large subunit (TerL), which is a powerful motor used to package DNA into the phage capsid. We have previously used terminases as phage markers to study phage-host interactions within the termite gut (70). Unlike many other viral genes such as integrases and lysins, terminases lack bacterial homologs, and thus, are considered to be unique to viruses (51, 52). In meeting the second criterion, pairwise sequence similarity analysis was performed to exclude candidate markers that shared any significant similarity. In exploring thousands of double-stranded DNA phage genomes, we have found that terminases from different phages typically do not share any significant sequence similarity (55). In cases where we detected sequence similarity between two terminase sequences, they belonged to highly similar phages infecting the same host species (55). As such, by imposing the second criterion, we likely arrived at distinct TerL lineages, with each lineage representing a distinct phage family.

Moreover, to meet the third criterion, we used two small oral metagenomic datasets (59, 71) and chose only TerL sequences that appeared in more than one individual. Together, these criteria led us to 7 non-homologous TerL lineages. We then searched for these 7 markers across larger, publically available metagenomic datasets (24, 72), and found the markers to be present across many individuals (Tadmor *et al*., in preparation). Now that we had used shotgun metagenomic datasets to identify several ubiquitous phage markers, we aimed to use the benefits of targeted sequencing to develop a high-resolution survey of phage communities across space and time.

### The sample collection kit and measures against sampling contamination

To obtain samples, we developed a sample collection kit and prepared kit contents within the PCR flowhood. Before and after every kit preparation session, the flowhood surfaces and pipettes were wiped using sterile wipes, DNA AWAY™, and 95% ethanol. At the end of each session the surfaces were also UV-sterilized (60 minutes). Each kit contains plastic tongue scrapers (Yellow CeraSpoon Safe Ear Curettes, Bionix) that were first autoclaved and then UV-sterilized for 60 minutes, 1.5 mL gamma-sterilized and pre-packaged collection tubes certified as pyrogen-RNase-DNA-and ATP-free (VWR), each containing 200 uL sterile 1X PBS buffer (VWR), along with pre-packaged sterile gloves (VWR). Each collection tube and tongue scraper pair was placed inside a sterile bag and the bags were placed in another bag. The next steps were performed outside of the flowhood. Each collection bag was put inside a Styrofoam box along with ice gel packs. Ice gel packs and Styrofoam boxes were not reused to prevent cross contamination between individuals in case of a spill, which would already be highly unlikely due to multiple layers of packaging. Upon arrival of samples, collection tubes were taken out of their original bags, wiped with 95% ethanol and DNA AWAY™ using sterile wipes and placed into a new sterile bag. Gloves were frequently exchanged both during this step and before proceeding to the next collection tube to prevent cross contamination. In addition to standard lab attire such as gloves and lab coat, a facemask was worn to prevent contamination during kit preparation and sample storage.

### Subject recruitment and sample collection

For the bulk of our sample collection, we relied heavily on citizen scientists. We made an educational video to introduce a diverse audience to the fascinating world of phages, explain our study and to recruit volunteers. We also created an instructional video for prospective volunteers on subject disqualifying criteria and subject rights, and to provide a step-by-step demonstration of sample collection, storage, and shipment. Among other exclusion criteria, subjects could not have taken antibiotics for the preceding 3 months and subjects could not have active cavities or gum disease. Qualified subjects were sent a kit and were asked not to brush their teeth or tongue for a minimum of 8 hours prior to sample collection to allow for a substantial build up of plaque on the tongue dorsum. Put simply, subjects were instructed to 1) wear gloves, 2) scrape their tongue (dorsal surface) several times using the tongue scraper, 3) deposit their sample into the collection tube, 4) place the tube back into the bag, and 5) store the bag in their freezer along with ice gel packs prior to an over-night shipment of their samples. They were also instructed to report any sources of error that occurred at any step, and to send their samples along with their signed consent form and questionnaire. Our sample collection and processing protocols were approved by Caltech Institutional Review Board (IRB protocol 14-0430) and Institutional Biosafety Committee (IBC protocol 13-198).

Nine subjects included in this study are those included in a previous study of oral microbial diversity by Bik *et al*. (21). Briefly, samples were collected from individuals by a dentist who examined subjects for their oral health, thereby excluding subjects with active cavities, gingivitis, or periodontal disease. For each subject, samples from different oral sites were collected using sterile curettes and deposited separately in 1.5 mL collection tubes containing PBS buffer. The 6 oral sites sampled include plaque from tongue dorsum, tongue ventral, buccal mucosa, hard palate, supra-gingiva, and sub-gingiva.

### Measures against contamination

A common source of contamination in PCR originates from previously amplified template sequences that enter new PCR reactions. To prevent contamination this type of contamination, four physically separated workstations were developed for DNA extraction (station A1), PCR preparation (station A2), PCR and gel electrophoresis (station B1), and PCR cleanup (station B2). A and B specify two different buildings at Caltech while 1 and 2 refer to two different rooms within the same building. The flow of materials was from building A to B and never the vice-versa. Every station had its own set of lab equipment, materials, and storage space. Disposable lab coats (Sigma-Aldrich^®^) were worn and disposed of at the end of every procedure to ensure that DNA was not carried between stations via clothing. Facemasks (Fisher Scientific) were also worn at all times to prevent any oral or nasal droplets from entering reactions. Prior to the start of every DNA extraction, lab equipment and bench tops were cleaned using sterile wipes and DNA AWAY™ (Thermo Scientific), a surface decontaminant that eliminates DNA and DNAses. PCR preparations and aliquoting of reagents were carried out in a PCR flowhood (AirClean^®^ Systems) equipped with a UV light and laminar airflow capabilities. Lab equipment required for PCR preparation was designated to the PCR preparation flowhood. At the end of every experimental session and when introducing new equipment into the flowhood, all surfaces were first wiped with DNA AWAY™ solution and then exposed to UV radiation for 60 minutes. Prepackaged, sterile gloves were used for PCR preparation. To prevent sample-to-sample contamination during DNA extraction, PCR preparation, and PCR cleanup, gloves were frequently exchanged. Most importantly, 5 No Template Control (NTC) reactions accompanied every PCR run. Similarly, to test the presence of contaminants in extraction reagents, for every extraction experiment, 3 reactions were carried out without the addition of any sample. PCR using phage primers was performed on these extraction control reactions.

### DNA Extraction (Station A1)

DNA extraction of human oral samples was done according to the manual from MoBio PowerBiofilm^®^ DNA Isolation Kit. The advantage to using this kit for DNA extraction and purification is that it combines the use of chemical and mechanical (bead-beating) treatments for an increased efficiency in biofilm disruption, lysis, and removal of inhibitors such as humic acid. The final concentrations of DNA were measured using Nanodrop. The concentration range of the total extracted genomic DNA was typically between 5 to 50 ng/µL.

### PCR preparation (Station A2) and PCR (Station B1)

Each PCR reaction contained 12.5 µL of PerfeCTa^®^ qPCR SuperMix, ROX™ (Quanta Biosciences), a premix containing AccuStart™ Taq DNA polymerase, MgCl_2_, dNTPs, and ROX reference dye for qPCR applications. Additionally, each reaction contained 10.5 µL of RT-PCR Grade Water (Ambion^®^) which is free of nucleic acids and nucleases, 1 µL of extracted DNA at 1 ng/µL, 0.5 µL of forward and 0.5 µL of reverse primers, each at 50 ng/µL (synthesized by IDT). A higher than recommended primer concentration was used because the phage primers used are 32-64 fold degenerate. The thermocycling protocol was made according to PerfeCTa qPCR SuperMix recommendations: 1) a 10-minute activation of AccuStart™ Taq DNA polymerase at 95°C, 2) 10 seconds of DNA denaturation at 95°C, 3) 20 seconds of annealing at 60°C, and 4) 30 seconds of extension at 72°C, 40 cycles repeating steps 2 to 4, followed by 5 minutes of final extension at 72°C.

### Gel electrophoresis (Station B1) and PCR cleanup (Station B2)

Phage PCR products were visualized using 2% agarose in TAE buffer. After gels were cast, 5 µL of each PCR product was mixed with 1 µL of 6X loading dye and loaded into a well. 5 µL of 100 base-pair ladder was used, and the gel electrophoresis instrument was set to run for 30 minutes at 100V. Phage PCR positive hits were purified using the QIAquick PCR Purification Kit (QIAGEN). 20µL of PCR products were used and purified according to the QIAquick PCR Purification manual.

### Illumina sequencing

Upon PCR cleanup, double stranded DNA concentration in each sample was measured using Qubit instrument. Qubit measurements were performed in Building C due to practical considerations rather than a necessary treatment for preventing contamination. Samples were combined into one reaction (∼2 µg dsDNA) and submitted to GENEWIZ, Inc for library preparation and MiSeq 2×300bp Paired-End sequencing.

### DNA barcodes for multiplexed sequencing

To enable multiplexing, unique DNA barcodes (Table 1) were appended onto the forward primer sequences (Table 3) used to amplify each phage marker. These barcoded primer sequences were synthesized by IDT. Using this scheme, ∼100 samples were submitted per MiSeq sequencing run (Table 1) and by matching the barcode sequence to the sample ID, information about who and where the sample came from was accessible. More specifically, Hamady error-correcting 8-letter barcodes (73) were used. Hamady DNA barcodes are an example of Hamming code wherein the addition of parity bits allow for detection and correction of errors within the barcode sequence. In the case of Hamady barcodes, up to 2 errors in the barcode sequence can be detected and one error can be corrected.

### Quality control steps to eliminate sequencing errors

We used Illumina MiSeq’s 2×300bp paired-end configuration (GENEWIZ, Inc). Each sequencing run produced about 20-25 Million paired-end reads. Paired-end reads were joined using *join_paired_ends.py* script from QIIME (Quantitative Insights Into Microbial Ecology) package, and unless noted otherwise scripts used in this chapter are part of QIIME (74). When a base is confirmed by both reads, higher Phred score is increased by up to 3 points. If paired reads had any mismatches across their overlapping bases, the paired reads was eliminated from any further analysis (QC step #1). For markers HB1, PCA2, and HA the overlap between the paired reads entirely covers the marker sequence, hence eliminating many sequencing errors.

Upon joining reads and eliminating those with mismatches in the region of overlap *seqQualityFilters.py*, an in-house script, was used to preform QC step #2: taking joined reads from QC step #1, and eliminating any sequences that have one or more bases marked by a Phred score below 30. Excluded from QC step #2 were the first two bases in the beginning and end of each sequence, which for majority of reads have much lower quality scores.

Using *seqQualityFilters.py,* sequences were placed in 3 different bins according to their primer sequences, and any sequence that did not have the correct barcode length, or the correct primer sequences at the expected positions, was eliminated (QC step #3). Additionally, nearly half of remaining sequences had to be reverse complemented so that all sequences were oriented in the 5’ to 3’ direction. Using the same script, primer and barcode sequences were removed, and barcode sequences were written to a separate file (to be used as input to *split_libraries_fastq.py*). At this point sequences that did not have the correct length were filtered out (QC step #3). Sequences were demultiplexd using *split_libraries_fastq.py* and reads with errors in the barcode sequence were eliminated (QC step #4).

### Phage community composition plots (“Phageprints”)

After demultiplexing quality-controlled reads, sequences were clustered according to a specified sequence similarity threshold using UCLUST *de novo* clustering algorithm (75) used in *pick_otus.py* script. Using *make_otu_table.py*, OTU tables were generated. An OTU table summarizes counts of sequences assigned to each OUT across each sample. We refer to this per-sample sequence count as the OTU size. As long as an OTU of size 1 or greater exists in at least one sample, it is included in the OTU table. In this way, the counts of OTUs for samples containing the same marker remains the same, though their size could vary widely across different samples. Later we will demonstrate the effects of noise filters applied to the OTU table. The relative abundance of each OTU within each sample was calculated via *processOtuTable.py,* another in-house script. In plotting the relative OTU abundance values for different samples, we arrived at complex, individual-specific patterns. We dubbed these phage community composition plots as “phageprints”.

### Metrics for quantitative comparison of phageprints

The first metric explored is binary Jaccard distance, which is equal to one minus the ratio of the intersection to the union of two samples’ OTUs: 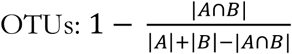. Here, *A* and *B* represent the OTUs that are present in sample 1 and 2, respectively. This is a binary method of comparing samples simply based on the presence/absence of the OTUs. In addition to the Pearson distance (1-Pearson correlation), we chose two other abundance-based distance metrics, namely abundance-weighted Jaccard and Bray-Curtis. Abundance-weighted Jaccard, which is equal to 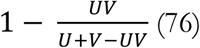 (76), is similar to Jaccard but here *U* and *V* represent the sum of relative abundances of OTUs shared between samples 1 and 2, respectively. Bray-Curtis dissimilarity (77) is defined as 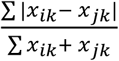, where *x*_*ik*_ and *x*_*jk*_ correspond to the relative abundance of OTU *k* in samples *i* and *j.*

Lastly, we explored unweighted Unifrac, a phylogenetic distance metric (78). The Unifrac algorithm operates on a phylogenetic tree containing sequences from all samples. It proceeds to create pairwise comparisons between samples by identifying the branch lengths that are shared between two samples, as well as the branch lengths that are unique to each sample. The Unifrac distance is then defined as the unshared branch lengths divided by the total branch lengths, where total branch lengths is the sum of shared and unshared branch lengths. If two samples are identical, the fraction of the tree’s branch lengths that is unique to one sample or the other will be zero, and thus, the Unifrac distance will be zero.

### Examining the effect of OTU sequence similarity threshold

In analyzing 16S sequences, clusters or Operational Taxonomic Units (OTUs) are conventionally defined at 97% sequence similarity threshold. To examine the effect of sequence similarity threshold for phage OTU formation, we tested OTU sequence similarity thresholds of 98%, 97%, 95%, 90%, and 80%. Figure 14 is a matrix of Pearson correlation coefficients calculated during the pairwise comparison of HB1 community compositions using different sequence similarity thresholds for defining OTUs. Very similar Pearson correlation matrices are obtained as the sequence similarity threshold is lowered from 98% to 80%. However, because the number of cluster is reduced as we reduce the sequence similarity threshold, with lower sequence similarity thresholds, the chance that individual-specific variations are lumped into the same cluster is increased. If noise-induced sequence variations are effectively accounted for, higher sequence similarity thresholds for defining OTUs can enable a more accurate and detailed depiction of a person’s phage community composition. For this reason, we used a sequence similarity threshold of 98% for the study of different oral sites, and later we used a 100% sequence similarity threshold for the temporal and the global study.

**Figure 13.**
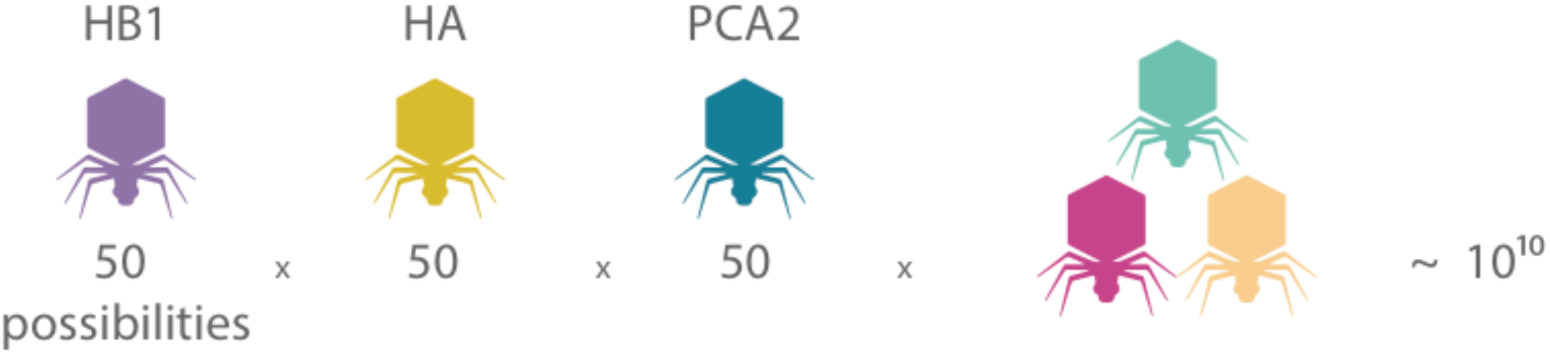
An estimate for the number of additional globally-distributed phage families needed to achieve the number of possible phageprint patterns that surpass the current human population. Assuming that phageprints from each phage family can provide 50 unique patterns, there would only be 3 additional globally distributed phage families needed.

**Figure 14.**
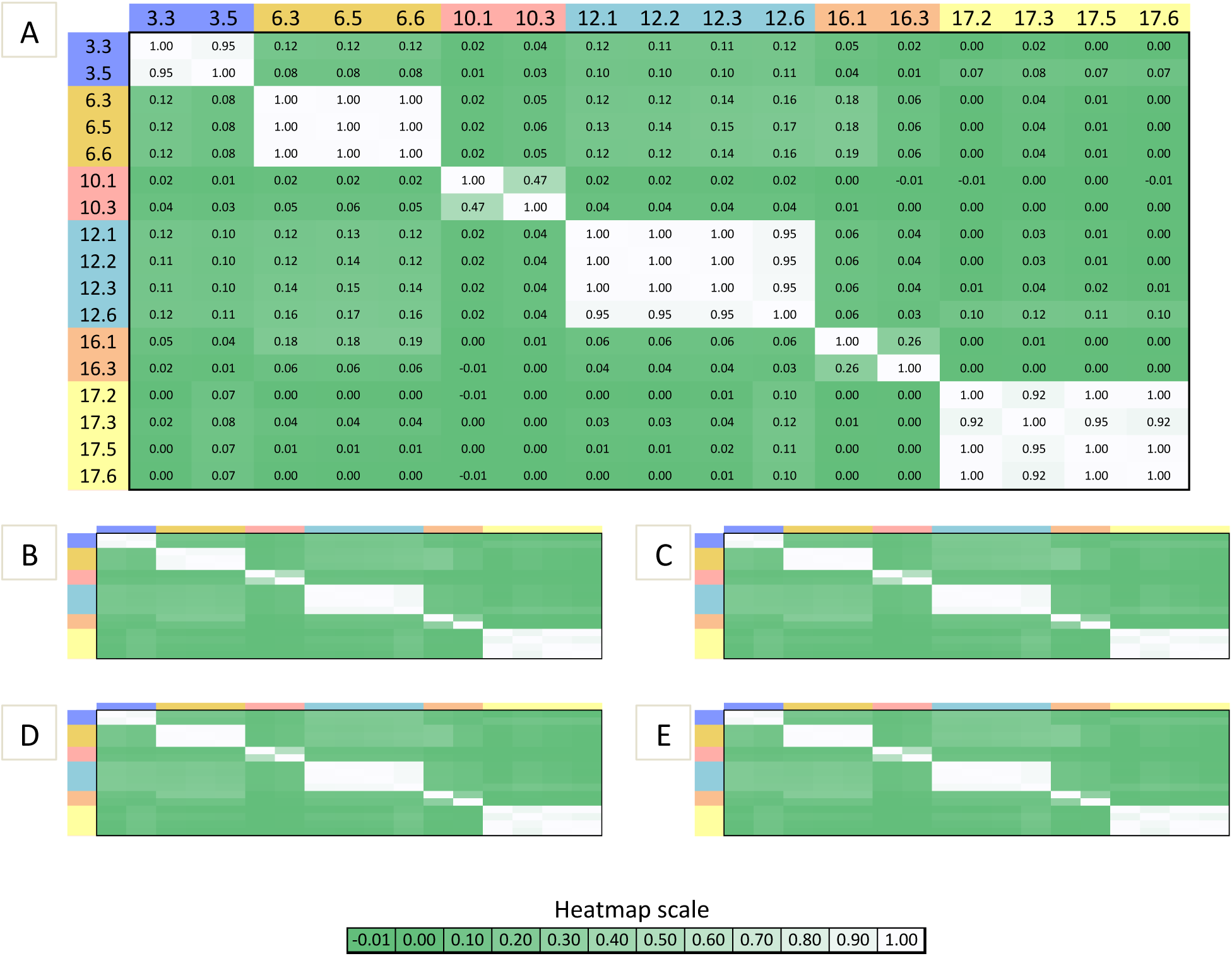
Pairwise Pearson correlation coefficient values calculated for HB1 phage community compositions as a function of A) 98%, B) 97%, C) 95%, D) 90%, and E) 80% sequence similarity thresholds for OTU formation. Sample IDs can be decoded as before: subject ID precedes oral site ID. Oral sites 1-6 correspond to tongue dorsum, hard palate, buccal mucosa, ventral tongue, supra-gingiva, and sub-gingiva respectively (e.g. 3.3 corresponds to subject 3 community composition derived from the buccal mucosa, and 3.5 is subject 3 supra-gingiva community composition). The number of OTUs generated at 98%, 97%, 95%, 90%, and 80% sequence similarity thresholds are 210, 181, 172, 170, and 80, respectively.

### Detecting experimental noise

How reproducible is a phage community composition plot? Figure 15 summarizes the sources of noise from all experimental processes performed during this study. First, it’s important to capture sampling variation. How consistently can we capture a phage community from an individual’s oral site given that we are sampling different parts of the biofilm each time? Another factor that could contribute to sampling variation are the personal differences in the rate of biofilm mass accumulation on the tongue dorsum. Secondly, we need to ask whether processes of lysis and DNA extraction allow for the availability of the same template DNA sequences in the same relative abundances across different extraction runs.

**Figure 15.**
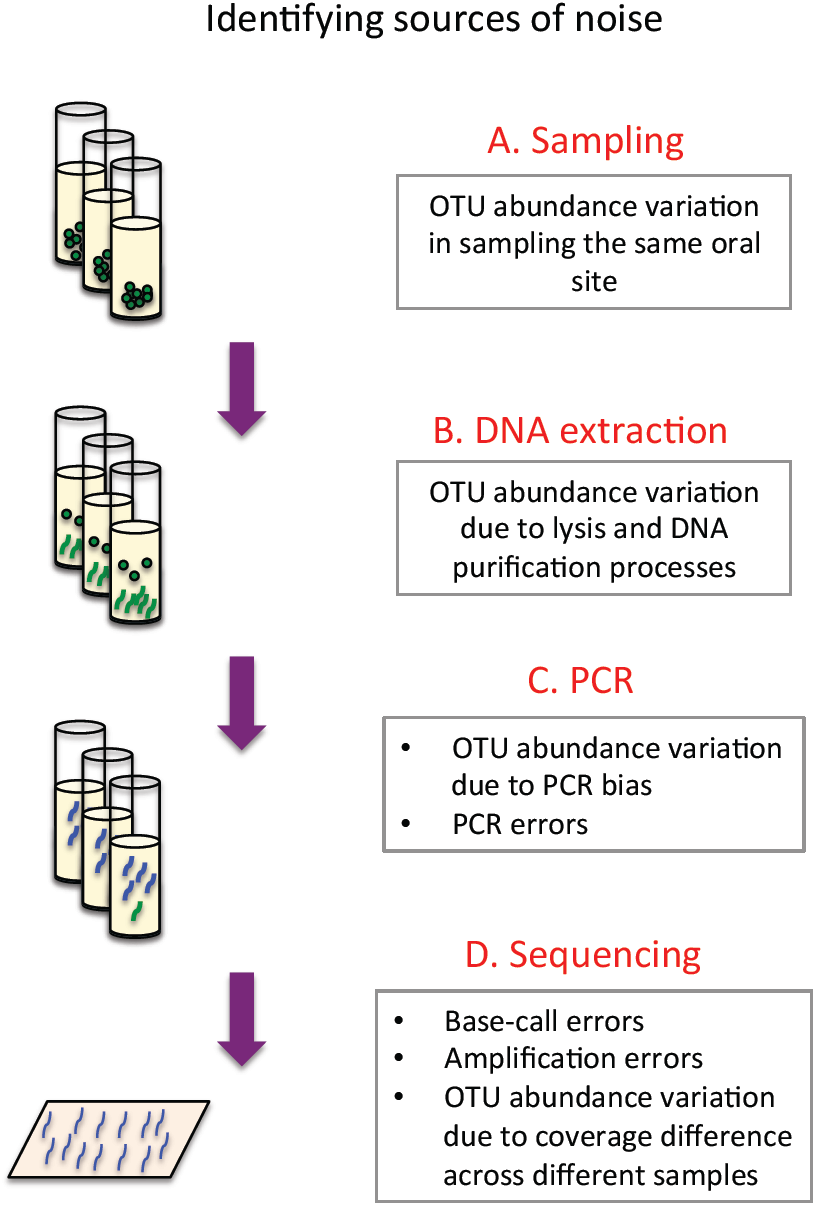
Sources of error and variation in experimental processes used in this study. A) Sampling of the same oral site in the same individual could result in collection of different microbial communities, which could introduce new OTUs or change relative abundance of existing OTUs. B) DNA extraction is not 100% efficient and the fraction of DNA extracted from an environment could serve as a source of variation across different samples. C) PCR introduces errors that could present themselves as novel OTUs or cause variation in abundance of genuine OTUs. D) Sequencing also introduces errors both at the level of base-calling and bridge amplification.

Third, we need to evaluate the OTU abundance variations that could result in PCR due to both errors as well as other stochastic events. For example, it’s possible that very rare template sequences are left out of the initial cycles of PCR and their relative abundance at the end of PCR is lower than their relative abundance prior to PCR. In this hypothetical scenario PCR could serve as a biased amplifier. PCR purification is similar to extraction and sampling in that it does not introduce sequence errors; however it is unlike these processes because after PCR billions of template copies are created and it’s unlikely that the loss of a fraction of templates during PCR purification will dramatically change OTU relative abundances. Finally, Illumina MiSeq sequencing is another error-prone process not only at the level of base-calling, but at the level of bridge amplification which like PCR could introduce errors that propagate exponentially. Refer to Figure 15 for a summary of processes that could result in irreproducible OTUs or variation in OTU relative abundances.

To quantify how reproducible a given phage community composition is, we obtained 3 different samples from subject 37 tongue dorsum. We then performed DNA extraction and PCR separately on each sample and sent samples for sequencing (sequencing run #2). The logic behind this experiment was to capture a lumped measure of noise arising from various processes depicted in Figure 15. After performing quality control steps 1-4, demultiplexing reads based on their barcode sequences, clustering reads based on 98% sequence similarity threshold for OTU formation, rarefying the OTU table to 4000 reads per sample, and calculating the relative abundances of OTUs, we measured the standard deviation in the relative abundance of each OTU across these three samples (Figure 16). Remarkably, relative abundance values across these three samples were highly consistent, with the majority of OTUs having standard deviations below 0.2% and the maximum standard deviation observed was less than 0.7% relative abundance.

**Figure 16.**
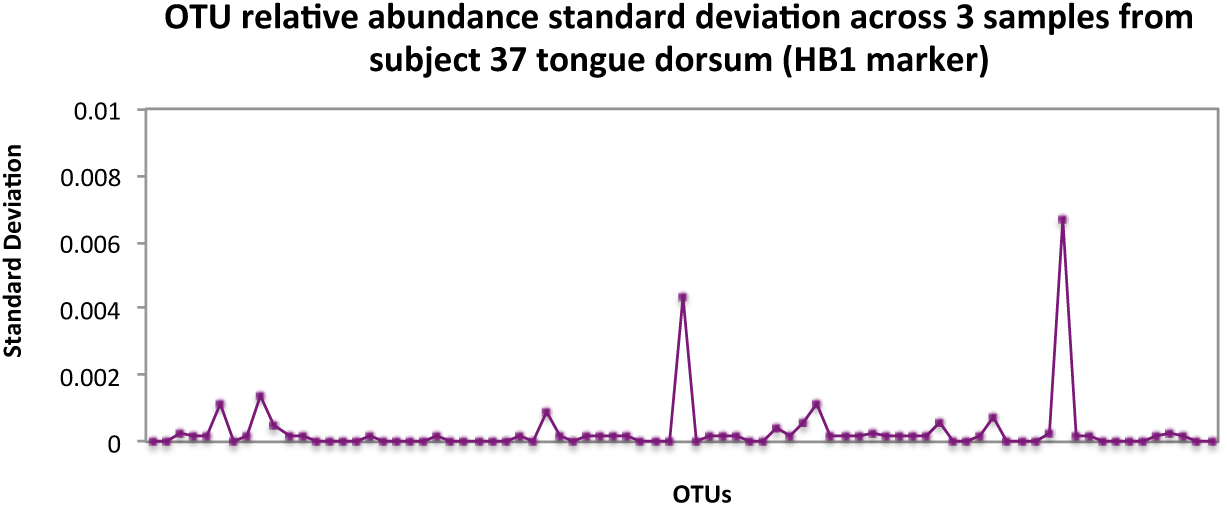
Standard deviations of OTU relative abundances calculated for all experimental processes. Three data points per OTU are used for standard deviation calculations. These three data points correspond to measurements of OTU relative abundances obtained for three different samples obtained from subject 37 tongue dorsum (HB1 marker) which underwent separate sampling, DNA extraction, PCR and PCR cleanup procedures. The maximum standard deviation observed is less than 0.007 relative abundance, and majority are close to 0.

### Identifying non-reproducible OTUs

To identify OTUs that were non-reproducible across the three samples from subject 37’s tongue dorsum (HB1 marker), we flagged OTUs that had appeared in only one or two samples out of three. We then plotted the histogram of non-reproducible OTUs as a function of their relative abundance (for those OTUs appearing in 2 out 3 samples, the higher relative abundance value was used). The thresholds defining each bin, *b*, were selected to be the following: 0>b_1_≥0.00025 (OTU of size 1 sequence since the total number of sequences per sample is 4000), 0.00025>b_2_≥0.0005 (2 sequences), 0.0005>b_3_≥0.00075 (3 sequences), 0.00075>b_4_≥0.001 (4 sequences), and 0.001<b_5_ (5 or more sequences).

Figure 17 demonstrates the number of non-reproducible OTUs drops as a function of OTU relative abundance, and all OTUs with more than 4 sequences (0.001 relative abundance) are reproducible. To conclude, we arrived at 0.001 relative abundance as the detection threshold for OTUs.

In addition to capturing a lumped sum of noise across all experimental processes for subject 37 tongue dorsum sample (Figure 16,Figure 17), for samples from subjects 3, 6, 10, 16, and 17, we performed a second set of PCR on previously extracted DNA samples, and submitted those samples for sequencing (Figure 18). In addition to these replicates, we acquired new samples from the tongue dorsum for subjects 31, 35, 37, and 38, and submitted these samples for the second sequencing run. In obtaining replicate phageprints, we were able to demonstrate that with proper quality filtration steps phageprints are highly reproducible even when they are generated from two separate PCR and sequencing steps (Figure 18).

**Figure 17.**
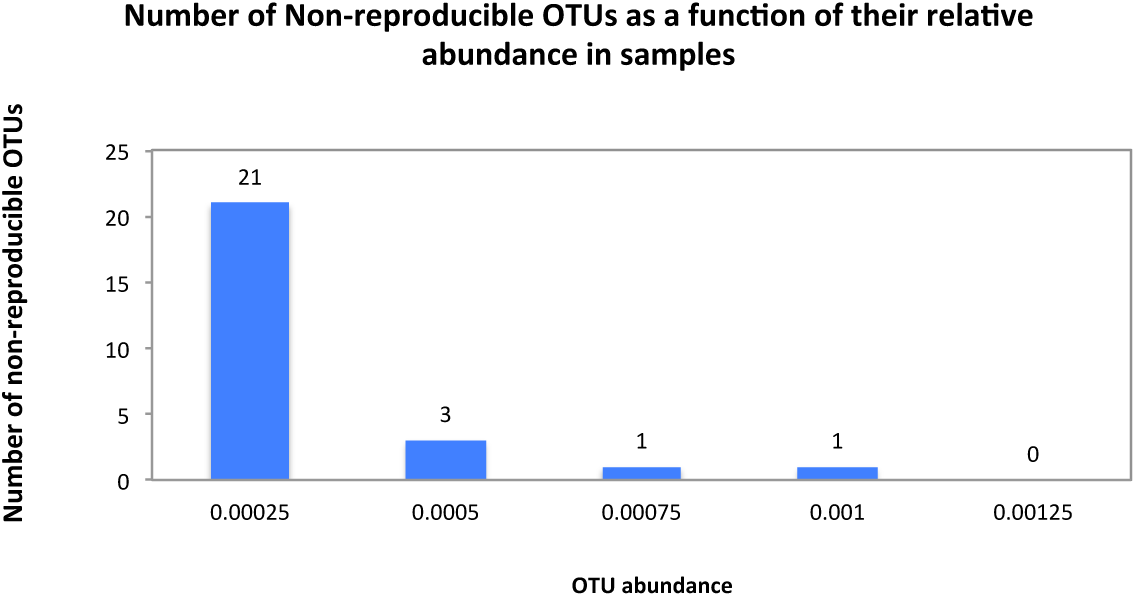
Number of non-reproducible OTUs across three samples obtained from subject 37 tongue dorsum (HB1 marker), presented as a function of OTU relative abundance. A total of 30 OTUs appear in one or two samples out of three, and therefore are considered non-reproducible. 21 out of 30 OTUs are defined by a single sequence which translates into 0.00025 relative abundance since samples are rarefied to 4000 sequences. The number of non-reproducible OTUs drops as a function of OTU relative abundance, and all OTUs with more than 4 sequences (0.001 relative abundance) are reproducible across three samples.

**Figure 18.**
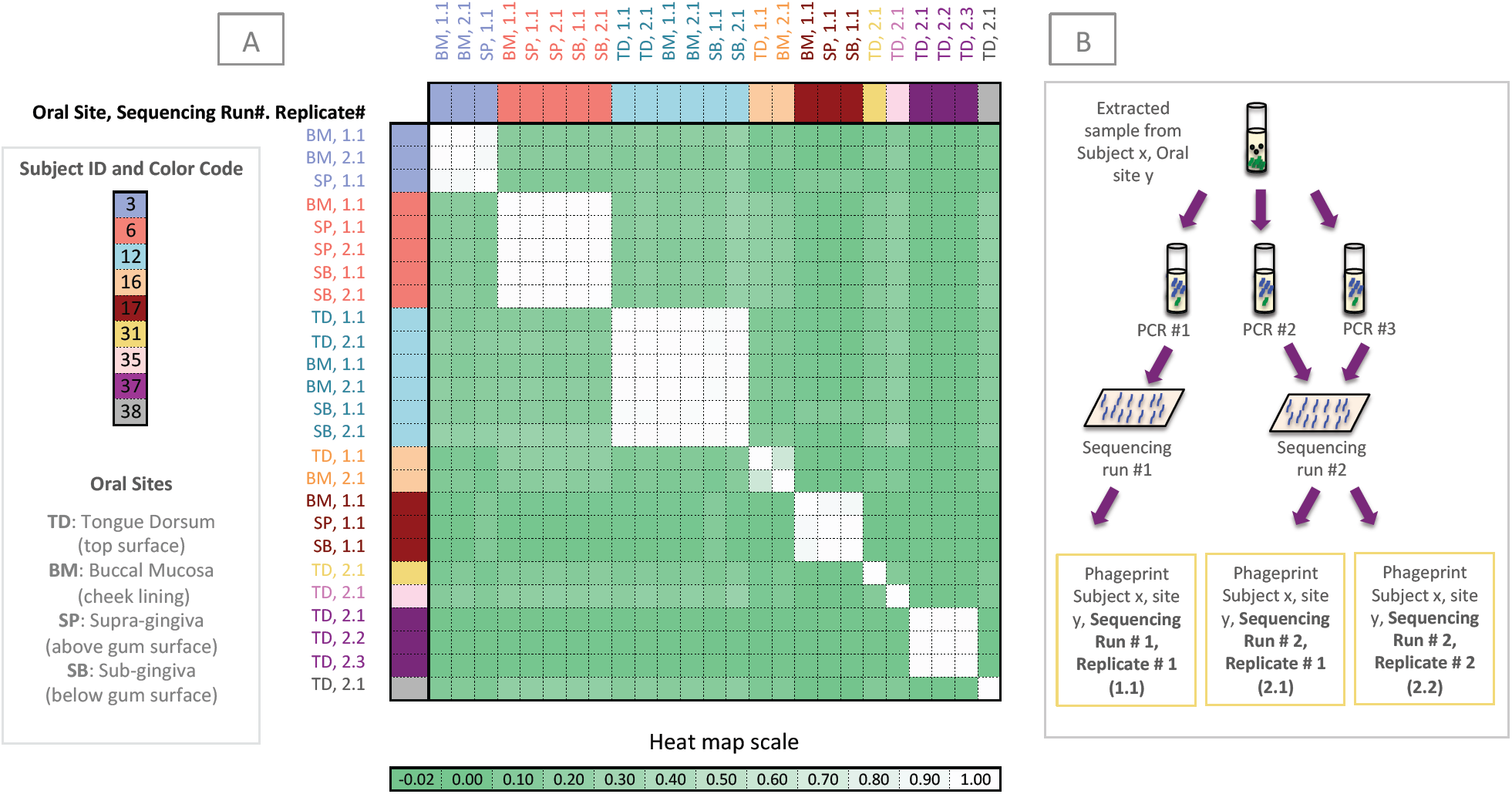
Panel A is the Pearson correlation matrix of all HB1 phageprints. Each phageprint is derived from the analysis of 4000 sequences associated with an individual and a particular oral site. OTUs are defined at 98% sequence similarity and OTUs with less than or equal to 0.1% relative abundance across all phageprints were filtered out. Phageprints are color-coded based on the individual they originate from. Oral sites shown to be positive for the HB1 marker are the tongue dorsum (TD), buccal mucosa (BM), supra-gingiva (SP), and sub-gingiva (SB). Phageprints that were acquired from sequencing run #1, are those marked as replicate #1. Panel B shows that to confirm reproducibility of phageprints, a second set of PCR was performed on previously extracted DNA from all samples included in sequencing run #1 and those PCR products were included in sequencing run #2. Phageprints derived from the second sequencing run are marked as replicate #2.

### Identifying phage marker homologs

The most abundant sequence from each OTU was retrieved using *pick_rep_set.py* to serve as a representative sequence. BLASTx function was used to detect the closest homolog to each OTU’s representative sequence from within the NCBI’s non-redundant protein database. HB1 representative sequences were aligned using Geneious (79), using a gap open penalty of 30 and gap extension penalty of 15 and a 65% similarity cost matrix. No gaps were introduced. The alignment is shown in SI Figure 4.

### Phage-host networks

OTU tables were input to *createNetwork.py,* an in-house script that creates node and edge tables. The nodes represent samples and phage OTUs, and a directed edge connects samples to the OTUs that they host. The weight of this connection is based on the relative abundance of the OTU in that sample. Gephi software (80) was used to visualize the resulting networks, and to obtain the degree distribution.

## Supplementary Information

**SI Figure 1.**
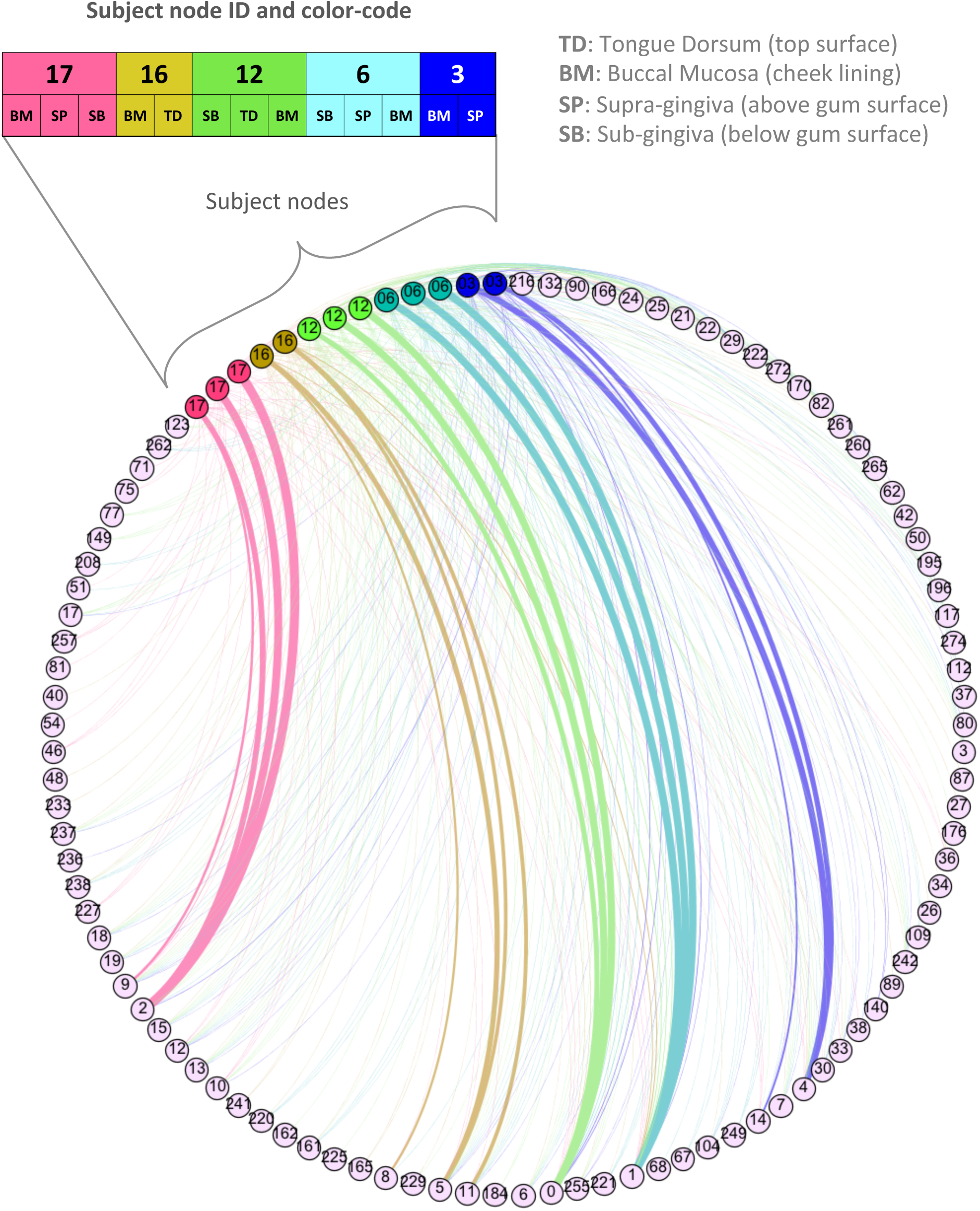
HB1 phage family network. Purple nodes are the OTU nodes and all other nodes represent samples. Sample nodes and edges are color-coded based on the individual they originate from. The oral site associated with each sample is abbreviated next to the sample’s node. Each edge connects an OTU a sample it exists in, and the edge weight is proportional to the relative abundance of the OTU in that sample. Node IDs are displayed. For OTU nodes, the node ID is the OTU ID which can be matched to IDs in SI Table 1 for identifying taxonomic information regarding each OTU. For sample nodes, the nodes IDs are simply the subjects’ IDs.

**SI Figure 2.**
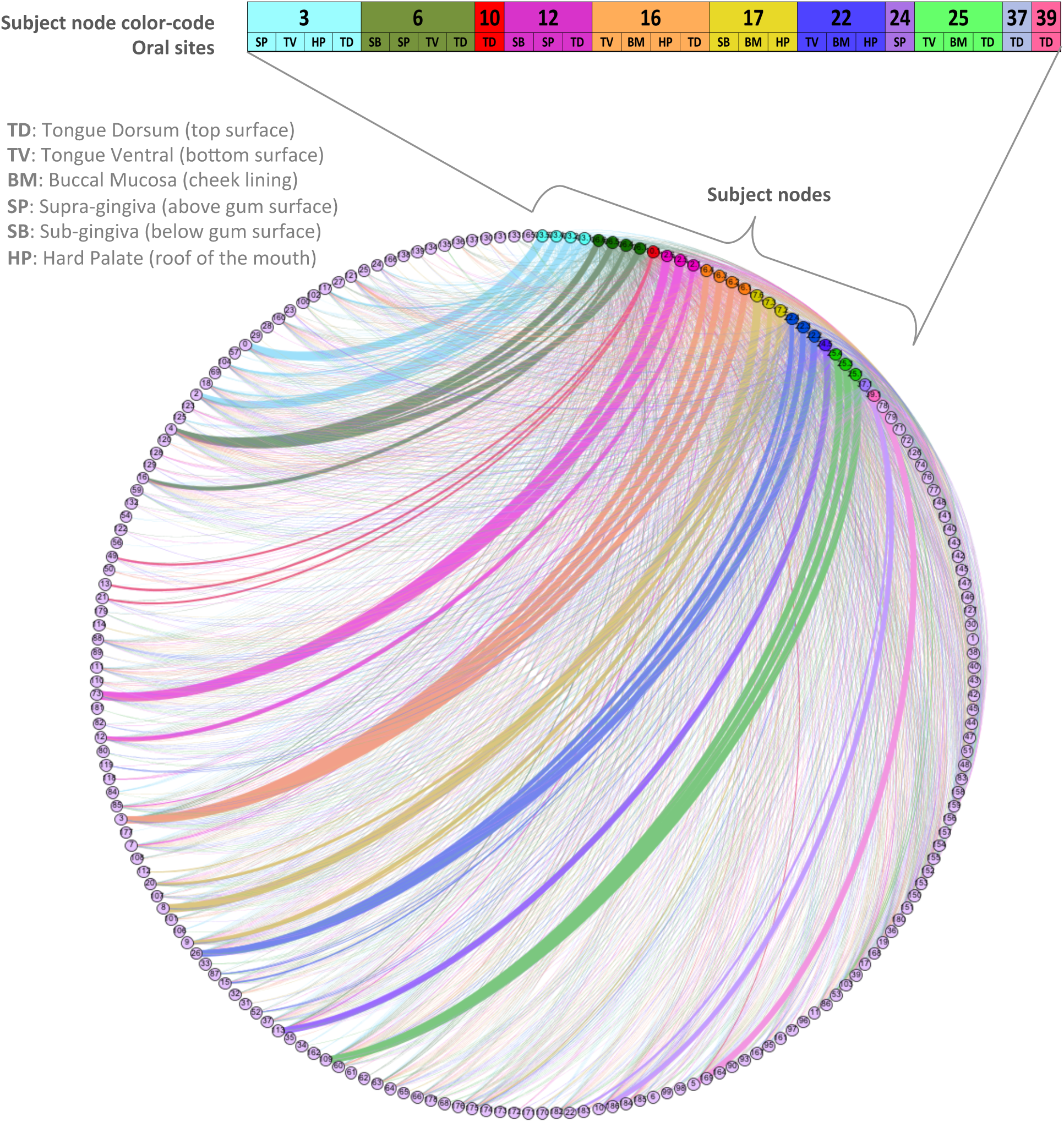
HA phage-host network. Purple nodes are the OTU nodes and all other nodes represent samples. Sample nodes and edges are color-coded based on the individual they originate from. Subject node color code, ID, and the oral sites are displayed above sample nodes. Each edge connects an OTU a sample it exists in, and the edge weight is proportional to the relative abundance of the OTU in that sample. Node IDs are displayed. For OTU nodes, the node ID is the OTU ID which can be matched to IDs in SI Table 3 for identifying taxonomic information regarding each OTU. For sample nodes, the nodes IDs are simply the subjects’ IDs.

**SI Table 1.** Closest homolog to each OTU’s representative sequence (HB1 phage family). Each OTU’s representative sequence was used as a query for NCBI’s BLASTx homology search against the non-redundant protein database. The table summarizes the E-value and the percent amino acid identity across the query sequence and the closest homolog, as well as the closest homolog’s name, sequence ID, and taxon ID. The taxon ID is color coded, and the taxonomic classification corresponding to each taxon ID can be retrieved from the following table. Note with the exception of a few “putative uncharacterized” homolog names that most are identified as terminases or TerLs (terminase large subunits).

| Query Sequence ID (OTU ID) | Percent Identity | E value | Closest Homolog | Closest Homolog Sequence ID | Closest Homolog Taxon ID |
| --- | --- | --- | --- | --- | --- |
| 0 | 71.6 | 2.00E-33 | TerL [Rhodococcus phage ReqiPepy6] | gi\|593779801\|ref\|YP_009017628.1\| | 691965 |
| 1 | 71.6 | 2.00E-33 | TerL [Rhodococcus phage ReqiPepy6] | gi\|593779801\|ref\|YP_009017628.1\| | 691965 |
| 10 | 67.9 | 3.00E-30 | terminase [[Clostridium] symbiosum] | gi\|489596073\|ref\|WP_003500516.1\| | 1512 |
| 103 | 69.14 | 5.00E-30 | terminase [[Clostridium] scindens] | gi\|639772655\|ref\|WP_024738760.1\| | 29347 |
| 104 | 71.6 | 2.00E-33 | TerL [Rhodococcus phage ReqiPepy6] | gi\|593779801\|ref\|YP_009017628.1\| | 691965 |
| 106 | 71.6 | 2.00E-33 | TerL [Rhodococcus phage ReqiPepy6] | gi\|593779801\|ref\|YP_009017628.1\| | 691965 |
| 109 | 71.6 | 2.00E-33 | TerL [Rhodococcus phage ReqiPepy6] | gi\|593779801\|ref\|YP_009017628.1\| | 691965 |
| 11 | 72.84 | 1.00E-34 | TerL [Rhodococcus phage ReqiPepy6] | gi\|593779801\|ref\|YP_009017628.1\| | 691965 |
| 112 | 71.6 | 2.00E-33 | TerL [Rhodococcus phage ReqiPepy6] | gi\|593779801\|ref\|YP_009017628.1\| | 691965 |
| 117 | 71.6 | 2.00E-33 | TerL [Rhodococcus phage ReqiPepy6] | gi\|593779801\|ref\|YP_009017628.1\| | 691965 |
| 118 | 71.6 | 2.00E-33 | TerL [Rhodococcus phage ReqiPepy6] | gi\|593779801\|ref\|YP_009017628.1\| | 691965 |
| 12 | 71.6 | 2.00E-33 | TerL [Rhodococcus phage ReqiPepy6] | gi\|593779801\|ref\|YP_009017628.1\| | 691965 |
| 122 | 70.37 | 2.00E-19 | TerL [Rhodococcus phage ReqiPepy6] | gi\|593779801\|ref\|YP_009017628.1\| | 691965 |
| 123 | 71.6 | 2.00E-33 | TerL [Rhodococcus phage ReqiPepy6] | gi\|593779801\|ref\|YP_009017628.1\| | 691965 |
| 128 | 71.6 | 2.00E-33 | TerL [Rhodococcus phage ReqiPepy6] | gi\|593779801\|ref\|YP_009017628.1\| | 691965 |
| 13 | 70.37 | 6.00E-33 | TerL [Rhodococcus phage ReqiPepy6] | gi\|593779801\|ref\|YP_009017628.1\| | 691965 |
| 132 | 70.37 | 4.00E-33 | TerL [Rhodococcus phage ReqiPepy6] | gi\|593779801\|ref\|YP_009017628.1\| | 691965 |
| 134 | 67.9 | 3.00E-30 | terminase [[Clostridium] symbiosum] | gi\|489596073\|ref\|WP_003500516.1\| | 1512 |
| 14 | 67.9 | 3.00E-30 | terminase [[Clostridium] symbiosum] | gi\|489596073\|ref\|WP_003500516.1\| | 1512 |
| 140 | 71.6 | 2.00E-33 | TerL [Rhodococcus phage ReqiPepy6] | gi\|593779801\|ref\|YP_009017628.1\| | 691965 |
| 142 | 71.6 | 2.00E-33 | TerL [Rhodococcus phage ReqiPepy6] | gi\|593779801\|ref\|YP_009017628.1\| | 691965 |
| 149 | 71.6 | 2.00E-33 | TerL [Rhodococcus phage ReqiPepy6] | gi\|593779801\|ref\|YP_009017628.1\| | 691965 |
| 15 | 71.6 | 5.00E-33 | TerL [Rhodococcus phage ReqiPepy6] | gi\|593779801\|ref\|YP_009017628.1\| | 691965 |
| 161 | 71.6 | 2.00E-33 | TerL [Rhodococcus phage ReqiPepy6] | gi\|593779801\|ref\|YP_009017628.1\| | 691965 |
| 162 | 71.6 | 2.00E-33 | TerL [Rhodococcus phage ReqiPepy6] | gi\|593779801\|ref\|YP_009017628.1\| | 691965 |
| 164 | 72.84 | 2.00E-32 | TerL [Rhodococcus phage ReqiPoco6] | gi\|593774729\|ref\|YP_009012597.1\| | 691964 |
| 165 | 71.6 | 2.00E-33 | TerL [Rhodococcus phage ReqiPepy6] | gi\|593779801\|ref\|YP_009017628.1\| | 691965 |
| 166 | 71.6 | 5.00E-33 | TerL [Rhodococcus phage ReqiPepy6] | gi\|593779801\|ref\|YP_009017628.1\| | 691965 |
| 17 | 71.6 | 2.00E-33 | TerL [Rhodococcus phage ReqiPepy6] | gi\|593779801\|ref\|YP_009017628.1\| | 691965 |
| 170 | 71.6 | 2.00E-33 | TerL [Rhodococcus phage ReqiPepy6] | gi\|593779801\|ref\|YP_009017628.1\| | 691965 |
| 176 | 71.6 | 3.00E-33 | TerL [Rhodococcus phage ReqiPepy6] | gi\|593779801\|ref\|YP_009017628.1\| | 691965 |
| 178 | 71.6 | 2.00E-33 | TerL [Rhodococcus phage ReqiPepy6] | gi\|593779801\|ref\|YP_009017628.1\| | 691965 |
| 18 | 71.6 | 2.00E-33 | TerL [Rhodococcus phage ReqiPepy6] | gi\|593779801\|ref\|YP_009017628.1\| | 691965 |
| 183 | 66.67 | 7.00E-30 | terminase [[Clostridium] symbiosum] | gi\|489596073\|ref\|WP_003500516.1\| | 1512 |
| 184 | 71.6 | 2.00E-33 | TerL [Rhodococcus phage ReqiPepy6] | gi\|593779801\|ref\|YP_009017628.1\| | 691965 |
| 189 | 71.6 | 4.00E-33 | TerL [Rhodococcus phage ReqiPoco6] | gi\|593774729\|ref\|YP_009012597.1\| | 691964 |
| 19 | 72.84 | 1.00E-34 | TerL [Rhodococcus phage ReqiPepy6] | gi\|593779801\|ref\|YP_009017628.1\| | 691965 |
| 195 | 71.6 | 2.00E-33 | TerL [Rhodococcus phage ReqiPepy6] | gi\|593779801\|ref\|YP_009017628.1\| | 691965 |
| 196 | 71.6 | 2.00E-33 | TerL [Rhodococcus phage ReqiPepy6] | gi\|593779801\|ref\|YP_009017628.1\| | 691965 |
| 197 | 71.6 | 2.00E-33 | TerL [Rhodococcus phage ReqiPepy6] | gi\|593779801\|ref\|YP_009017628.1\| | 691965 |
| 2 | 66.67 | 7.00E-30 | terminase [[Clostridium] symbiosum] | gi\|489596073\|ref\|WP_003500516.1\| | 1512 |
| 202 | 67.9 | 3.00E-30 | terminase [[Clostridium] symbiosum] | gi\|489596073\|ref\|WP_003500516.1\| | 1512 |
| 203 | 65.43 | 2.00E-28 | TerL [Rhodococcus phage ReqiPoco6] | gi\|593774729\|ref\|YP_009012597.1\| | 691964 |
| 206 | 70.37 | 2.00E-32 | TerL [Rhodococcus phage ReqiPepy6] | gi\|593779801\|ref\|YP_009017628.1\| | 691965 |
| 207 | 71.6 | 5.00E-33 | TerL [Rhodococcus phage ReqiPepy6] | gi\|593779801\|ref\|YP_009017628.1\| | 691965 |
| 208 | 71.6 | 2.00E-33 | TerL [Rhodococcus phage ReqiPepy6] | gi\|593779801\|ref\|YP_009017628.1\| | 691965 |
| 21 | 67.9 | 3.00E-30 | terminase [[Clostridium] symbiosum] | gi\|489596073\|ref\|WP_003500516.1\| | 1512 |
| 210 | 71.6 | 2.00E-31 | TerL [Rhodococcus phage ReqiPepy6] | gi\|593779801\|ref\|YP_009017628.1\| | 691965 |
| 213 | 69.14 | 2.00E-29 | TerL [Rhodococcus phage ReqiPoco6] | gi\|593774729\|ref\|YP_009012597.1\| | 691964 |
| 216 | 71.6 | 2.00E-33 | TerL [Rhodococcus phage ReqiPepy6] | gi\|593779801\|ref\|YP_009017628.1\| | 691965 |
| 218 | 71.6 | 2.00E-33 | TerL [Rhodococcus phage ReqiPepy6] | gi\|593779801\|ref\|YP_009017628.1\| | 691965 |
| 22 | 71.6 | 2.00E-33 | TerL [Rhodococcus phage ReqiPepy6] | gi\|593779801\|ref\|YP_009017628.1\| | 691965 |
| 220 | 71.6 | 2.00E-33 | TerL [Rhodococcus phage ReqiPepy6] | gi\|593779801\|ref\|YP_009017628.1\| | 691965 |
| 221 | 71.6 | 2.00E-33 | TerL [Rhodococcus phage ReqiPepy6] | gi\|593779801\|ref\|YP_009017628.1\| | 691965 |
| 222 | 71.6 | 2.00E-33 | TerL [Rhodococcus phage ReqiPepy6] | gi\|593779801\|ref\|YP_009017628.1\| | 691965 |
| 225 | 71.6 | 2.00E-33 | TerL [Rhodococcus phage ReqiPepy6] | gi\|593779801\|ref\|YP_009017628.1\| | 691965 |
| 227 | 66.67 | 7.00E-30 | terminase [[Clostridium] symbiosum] | gi\|489596073\|ref\|WP_003500516.1\| | 1512 |
| 229 | 71.6 | 2.00E-33 | TerL [Rhodococcus phage ReqiPepy6] | gi\|593779801\|ref\|YP_009017628.1\| | 691965 |
| 231 | 71.6 | 2.00E-33 | TerL [Rhodococcus phage ReqiPepy6] | gi\|593779801\|ref\|YP_009017628.1\| | 691965 |
| 233 | 71.6 | 2.00E-33 | TerL [Rhodococcus phage ReqiPepy6] | gi\|593779801\|ref\|YP_009017628.1\| | 691965 |
| 236 | 70.37 | 2.00E-32 | TerL [Rhodococcus phage ReqiPepy6] | gi\|593779801\|ref\|YP_009017628.1\| | 691965 |
| 237 | 71.6 | 2.00E-33 | TerL [Rhodococcus phage ReqiPepy6] | gi\|593779801\|ref\|YP_009017628.1\| | 691965 |
| 238 | 71.6 | 2.00E-33 | TerL [Rhodococcus phage ReqiPepy6] | gi\|593779801\|ref\|YP_009017628.1\| | 691965 |
| 24 | 71.6 | 5.00E-33 | TerL [Rhodococcus phage ReqiPepy6] | gi\|593779801\|ref\|YP_009017628.1\| | 691965 |
| 241 | 71.6 | 2.00E-33 | TerL [Rhodococcus phage ReqiPepy6] | gi\|593779801\|ref\|YP_009017628.1\| | 691965 |
| 242 | 71.6 | 2.00E-33 | TerL [Rhodococcus phage ReqiPepy6] | gi\|593779801\|ref\|YP_009017628.1\| | 691965 |
| 245 | 71.6 | 2.00E-33 | TerL [Rhodococcus phage ReqiPepy6] | gi\|593779801\|ref\|YP_009017628.1\| | 691965 |
| 249 | 71.6 | 2.00E-33 | TerL [Rhodococcus phage ReqiPepy6] | gi\|593779801\|ref\|YP_009017628.1\| | 691965 |
| 25 | 71.6 | 3.00E-33 | TerL [Rhodococcus phage ReqiPepy6] | gi\|593779801\|ref\|YP_009017628.1\| | 691965 |
| 250 | 71.6 | 2.00E-33 | TerL [Rhodococcus phage ReqiPepy6] | gi\|593779801\|ref\|YP_009017628.1\| | 691965 |
| 255 | 71.6 | 2.00E-33 | TerL [Rhodococcus phage ReqiPepy6] | gi\|593779801\|ref\|YP_009017628.1\| | 691965 |
| 256 | 72.84 | 4.00E-34 | TerL [Rhodococcus phage ReqiPepy6] | gi\|593779801\|ref\|YP_009017628.1\| | 691965 |
| 257 | 66.67 | 7.00E-30 | terminase [[Clostridium] symbiosum] | gi\|489596073\|ref\|WP_003500516.1\| | 1512 |
| 259 | 71.6 | 2.00E-33 | TerL [Rhodococcus phage ReqiPepy6] | gi\|593779801\|ref\|YP_009017628.1\| | 691965 |
| 26 | 71.6 | 5.00E-33 | TerL [Rhodococcus phage ReqiPepy6] | gi\|593779801\|ref\|YP_009017628.1\| | 691965 |
| 260 | 71.6 | 2.00E-33 | TerL [Rhodococcus phage ReqiPepy6] | gi\|593779801\|ref\|YP_009017628.1\| | 691965 |
| 261 | 71.6 | 2.00E-33 | TerL [Rhodococcus phage ReqiPepy6] | gi\|593779801\|ref\|YP_009017628.1\| | 691965 |
| 262 | 71.6 | 2.00E-33 | TerL [Rhodococcus phage ReqiPepy6] | gi\|593779801\|ref\|YP_009017628.1\| | 691965 |
| 264 | 71.6 | 3.00E-32 | TerL [Rhodococcus phage ReqiPoco6] | gi\|593774729\|ref\|YP_009012597.1\| | 691964 |
| 265 | 71.6 | 2.00E-33 | TerL [Rhodococcus phage ReqiPepy6] | gi\|593779801\|ref\|YP_009017628.1\| | 691965 |
| 27 | 62.96 | 2.00E-26 | TerL [Rhodococcus phage ReqiPepy6] | gi\|593779801\|ref\|YP_009017628.1\| | 691965 |
| 272 | 70.37 | 3.00E-32 | TerL [Rhodococcus phage ReqiPepy6] | gi\|593779801\|ref\|YP_009017628.1\| | 691965 |
| 273 | 67.9 | 2.00E-30 | TerL [Rhodococcus phage ReqiPoco6] | gi\|593774729\|ref\|YP_009012597.1\| | 691964 |
| 274 | 71.6 | 2.00E-33 | TerL [Rhodococcus phage ReqiPepy6] | gi\|593779801\|ref\|YP_009017628.1\| | 691965 |
| 29 | 74.07 | 2.00E-35 | TerL [Rhodococcus phage ReqiPepy6] | gi\|593779801\|ref\|YP_009017628.1\| | 691965 |
| 3 | 67.9 | 3.00E-29 | terminase [Clostridiales bacterium VE202-03] | gi\|639707411\|ref\|WP_024723669.1\| | 1232439 |
| 30 | 64.2 | 2.00E-28 | putative uncharacterized protein [Ruminococcus sp. CAG:17] | gi\|547240587\|ref\|WP_021976510.1\| | 1262951 |
| 32 | 67.9 | 3.00E-30 | terminase [[Clostridium] symbiosum] | gi\|489596073\|ref\|WP_003500516.1\| | 1512 |
| 33 | 67.9 | 3.00E-31 | TerL [Rhodococcus phage ReqiPoco6] | gi\|593774729\|ref\|YP_009012597.1\| | 691964 |
| 34 | 71.6 | 2.00E-33 | TerL [Rhodococcus phage ReqiPepy6] | gi\|593779801\|ref\|YP_009017628.1\| | 691965 |
| 36 | 76.54 | 2.00E-35 | TerL [Rhodococcus phage ReqiPepy6] | gi\|593779801\|ref\|YP_009017628.1\| | 691965 |
| 37 | 67.9 | 3.00E-30 | terminase [[Clostridium] symbiosum] | gi\|489596073\|ref\|WP_003500516.1\| | 1512 |
| 38 | 66.67 | 2.00E-29 | TerL [Rhodococcus phage ReqiPoco6] | gi\|593774729\|ref\|YP_009012597.1\| | 691964 |
| 4 | 71.6 | 2.00E-33 | TerL [Rhodococcus phage ReqiPepy6] | gi\|593779801\|ref\|YP_009017628.1\| | 691965 |
| 40 | 66.67 | 3.00E-32 | putative uncharacterized protein [Ruminococcus sp. CAG:17] | gi\|547240587\|ref\|WP_021976510.1\| | 1262951 |
| 42 | 64.2 | 3.00E-26 | terminase [[Clostridium] symbiosum] | gi\|489596073\|ref\|WP_003500516.1\| | 1512 |
| 44 | 66.67 | 5.00E-19 | terminase [[Ruminococcus] torques] | gi\|490985259\|ref\|WP_004846995.1\| | 33039 |
| 46 | 67.9 | 3.00E-30 | terminase [[Clostridium] symbiosum] | gi\|489596073\|ref\|WP_003500516.1\| | 1512 |
| 48 | 71.6 | 9.00E-34 | TerL [Rhodococcus phage ReqiPepy6] | gi\|593779801\|ref\|YP_009017628.1\| | 691965 |
| 5 | 71.6 | 2.00E-33 | TerL [Rhodococcus phage ReqiPepy6] | gi\|593779801\|ref\|YP_009017628.1\| | 691965 |
| 50 | 86.42 | 3.00E-44 | TerL [Rhodococcus phage ReqiPoco6] | gi\|593774729\|ref\|YP_009012597.1\| | 691964 |
| 51 | 67.9 | 3.00E-30 | terminase [[Clostridium] symbiosum] | gi\|489596073\|ref\|WP_003500516.1\| | 1512 |
| 54 | 65.43 | 6.00E-31 | terminase [[Clostridium] symbiosum] | gi\|489596073\|ref\|WP_003500516.1\| | 1512 |
| 59 | 71.6 | 2.00E-33 | TerL [Rhodococcus phage ReqiPepy6] | gi\|593779801\|ref\|YP_009017628.1\| | 691965 |
| 6 | 71.6 | 2.00E-33 | TerL [Rhodococcus phage ReqiPepy6] | gi\|593779801\|ref\|YP_009017628.1\| | 691965 |
| 60 | 71.6 | 2.00E-33 | TerL [Rhodococcus phage ReqiPepy6] | gi\|593779801\|ref\|YP_009017628.1\| | 691965 |
| 62 | 69.14 | 2.00E-30 | TerL [Rhodococcus phage ReqiPepy6] | gi\|593779801\|ref\|YP_009017628.1\| | 691965 |
| 67 | 71.6 | 2.00E-33 | TerL [Rhodococcus phage ReqiPepy6] | gi\|593779801\|ref\|YP_009017628.1\| | 691965 |
| 68 | 74.07 | 1.00E-34 | TerL [Rhodococcus phage ReqiPoco6] | gi\|593774729\|ref\|YP_009012597.1\| | 691964 |
| 7 | 71.6 | 2.00E-33 | TerL [Rhodococcus phage ReqiPepy6] | gi\|593779801\|ref\|YP_009017628.1\| | 691965 |
| 71 | 70.37 | 2.00E-31 | TerL [Rhodococcus phage ReqiPoco6] | gi\|593774729\|ref\|YP_009012597.1\| | 691964 |
| 72 | 62.96 | 4.00E-27 | TerL [Rhodococcus phage ReqiPoco6] | gi\|593774729\|ref\|YP_009012597.1\| | 691964 |
| 75 | 70.37 | 8.00E-33 | TerL [Rhodococcus phage ReqiPepy6] | gi\|593779801\|ref\|YP_009017628.1\| | 691965 |
| 77 | 71.6 | 2.00E-33 | TerL [Rhodococcus phage ReqiPepy6] | gi\|593779801\|ref\|YP_009017628.1\| | 691965 |
| 8 | 56.79 | 9.00E-26 | TerL [Rhodococcus phage ReqiPoco6] | gi\|593774729\|ref\|YP_009012597.1\| | 691964 |
| 80 | 69.14 | 3.00E-31 | TerL [Rhodococcus phage ReqiPoco6] | gi\|593774729\|ref\|YP_009012597.1\| | 691964 |
| 81 | 74.07 | 7.00E-33 | putative uncharacterized protein [Ruminococcus sp. CAG:17] | gi\|547240587\|ref\|WP_021976510.1\| | 1262951 |
| 82 | 71.6 | 2.00E-33 | TerL [Rhodococcus phage ReqiPepy6] | gi\|593779801\|ref\|YP_009017628.1\| | 691965 |
| 86 | 71.6 | 2.00E-33 | TerL [Rhodococcus phage ReqiPepy6] | gi\|593779801\|ref\|YP_009017628.1\| | 691965 |
| 87 | 67.9 | 1.00E-27 | terminase [[Clostridium] hathewayi] | gi\|493833739\|ref\|WP_006781000.1\| | 154046 |
| 89 | 74.07 | 1.00E-34 | TerL [Rhodococcus phage ReqiPoco6] | gi\|593774729\|ref\|YP_009012597.1\| | 691964 |
| 9 | 71.6 | 2.00E-33 | TerL [Rhodococcus phage ReqiPepy6] | gi\|593779801\|ref\|YP_009017628.1\| | 691965 |
| 90 | 71.6 | 2.00E-33 | TerL [Rhodococcus phage ReqiPepy6] | gi\|593779801\|ref\|YP_009017628.1\| | 691965 |

**SI Table 2.** Taxonomic classification of closest homologs (HB1 phage family). Majority of OTUs (86 out of 123) have the closest match to ReqiPoco6 terminase large subunit, whereas 15 OTUs have closest homologs belonging to ReqiPepy6.

| Closest<br>Homolog<br>Taxon ID | Kingdom | Phylum | Class | Order | Family | Genus | Species |
| --- | --- | --- | --- | --- | --- | --- | --- |
| 1262951 | Bacteria | Firmicutes | Clostridia | Clostridiales | Ruminococcaceae | Ruminococcus | Ruminococcus<br>sp. CAG:17 |
| 691964 | Viruses | dsDNA viruses,<br>no RNA stage | Caudovirales | Siphoviridae | unclassified<br>Siphoviridae | Rhodococcus<br>phage | ReqiPoco6 |
| 691965 | Viruses | dsDNA viruses,<br>no RNA stage | Caudovirales | Siphoviridae | unclassified<br>Siphoviridae | Rhodococcus<br>phage | ReqiPepy6 |
| 1512 | Bacteria | Firmicutes | Clostridia | Clostridiales | Lachnospiraceae | Lachnoclostridium | Clostridium<br>symbiosum |
| 29347 | Bacteria | Firmicutes | Clostridia | Clostridiales | Lachnospiraceae | Lachnoclostridium | Clostridium<br>scindens |
| 33039 | Bacteria | Firmicutes | Clostridia | Clostridiales | Lachnospiraceae | Blautia | Ruminococcus<br>torques |
| 1232439 | Bacteria | Firmicutes | Clostridia | Clostridiales | unclassified<br>Clostridiales | unclassified<br>Clostridiales | Clostridiales<br>bacterium VE202-03 |
| 154046 | Bacteria | Firmicutes | Clostridia | Clostridiales | Lachnospiraceae | Lachnoclostridium | Clostridium<br>hathewayi |

**SI Table 3.**
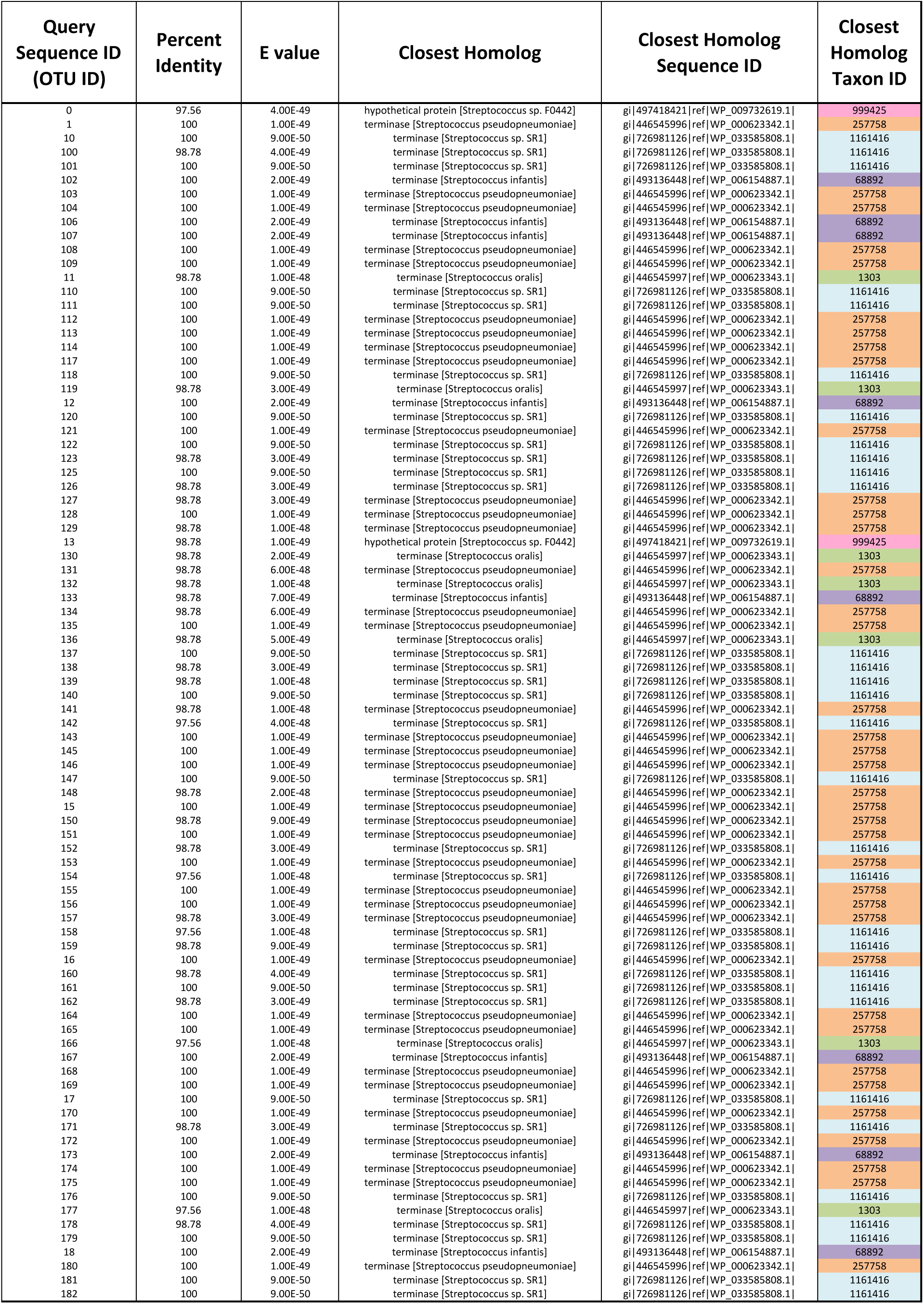
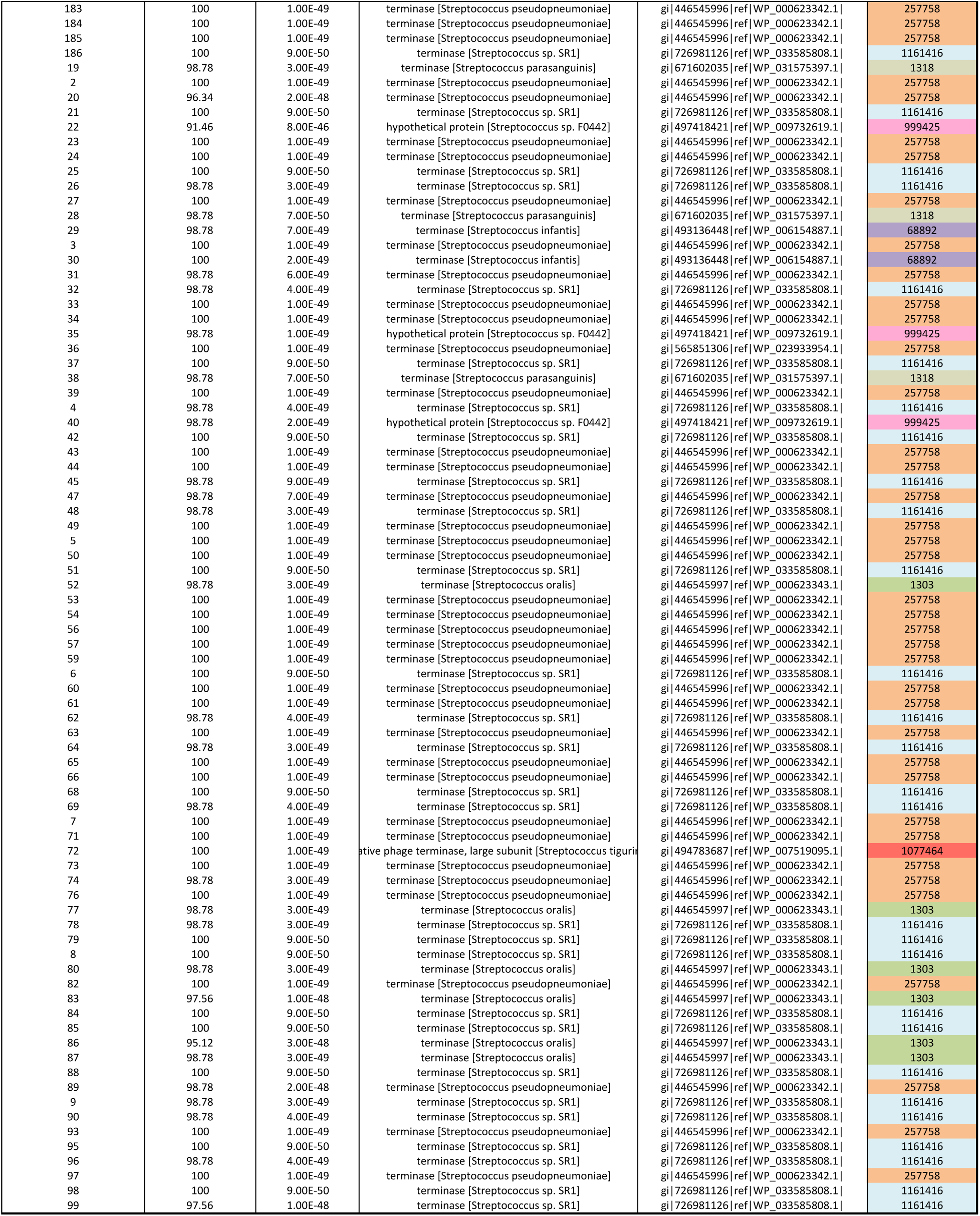
Closest homolog to each OTU’s representative sequence (HA phage family). Each OTU’s representative sequence was used as a query for NCBI’s BLASTx homology search against the non-redundant protein database. The table summarizes the E-value and the percent amino acid identity across the query sequence and the closest homolog, as well as the closest homolog’s name, sequence ID, and taxon ID. The taxon ID is color coded, and the taxonomic classification corresponding to each taxon ID can be retrieved from the following table. Note with the exception of a few “putative uncharacterized” homolog names that most are identified as terminases or TerLs (terminase large subunits).

**SI Table 4.** Taxonomic classification of closest homologs to each OTU’s representative sequence (HA phage family).

| Closest<br>Homolog<br>Taxon ID | Kingdom | Phylum | Class | Order | Family | Genus | Species |
| --- | --- | --- | --- | --- | --- | --- | --- |
| 1318 | Bacteria | Firmicutes | Bacilli | Lactobacillales | Streptococcaceae | Streptococcus | Streptococcus parasanguinis |
| 1077464 | Bacteria | Firmicutes | Bacilli | Lactobacillales | Streptococcaceae | Streptococcus | Streptococcus tigurinus |
| 257758 | Bacteria | Firmicutes | Bacilli | Lactobacillales | Streptococcaceae | Streptococcus | Streptococcus pseudopneumoniae |
| 999425 | Bacteria | Firmicutes | Bacilli | Lactobacillales | Streptococcaceae | Streptococcus | Streptococcus sp. F0442 |
| 1161416 | Bacteria | Firmicutes | Bacilli | Lactobacillales | Streptococcaceae | Streptococcus | Streptococcus sp. SR1 |
| 1303 | Bacteria | Firmicutes | Bacilli | Lactobacillales | Streptococcaceae | Streptococcus | Streptococcus oralis |
| 68892 | Bacteria | Firmicutes | Bacilli | Lactobacillales | Streptococcaceae | Streptococcus | Streptococcus infantis |

**SI Table 5.**
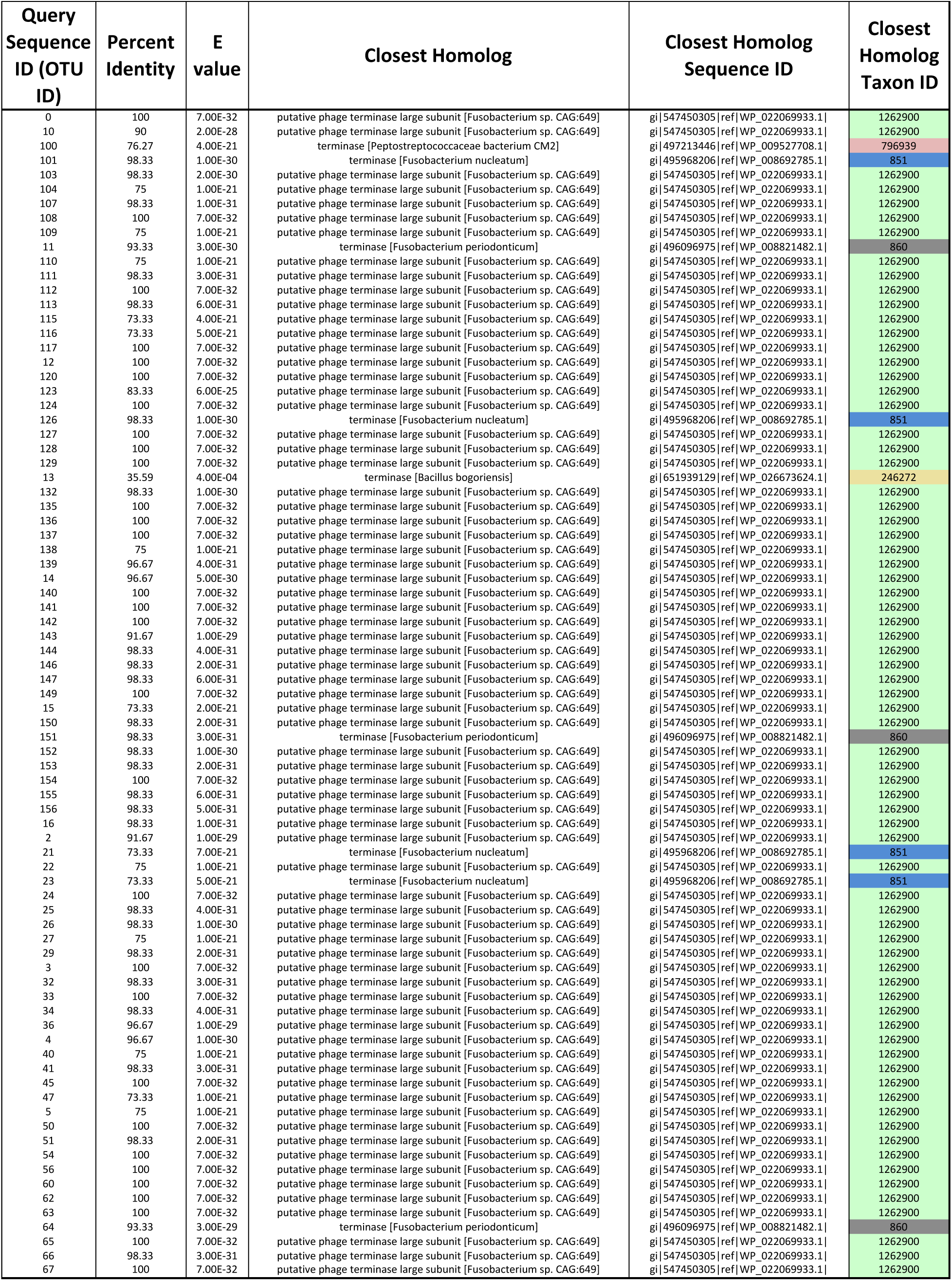
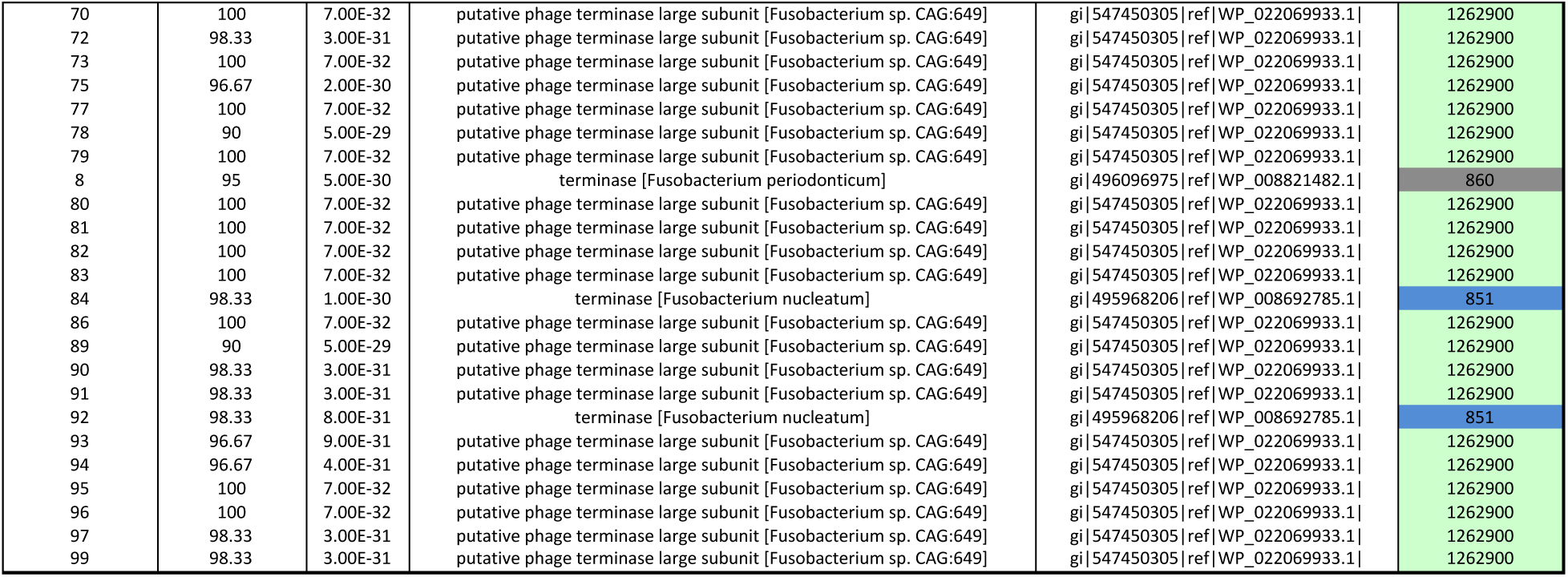
Closest homolog to each OTU’s representative sequence (PCA2 phage family). Each OTU’s representative sequence was used as a query for NCBI’s BLASTx homology search against the non-redundant protein database. The table summarizes the E-value and the percent amino acid identity across the query sequence and the closest homolog, as well as the closest homolog’s name, sequence ID, and taxon ID. The taxon ID is color coded, and the taxonomic classification corresponding to each taxon ID can be retrieved from the following table. Note with the exception of a few “putative uncharacterized” homolog names, most are identified as terminases or TerLs (terminase large subunits).

**SI Table 6.**
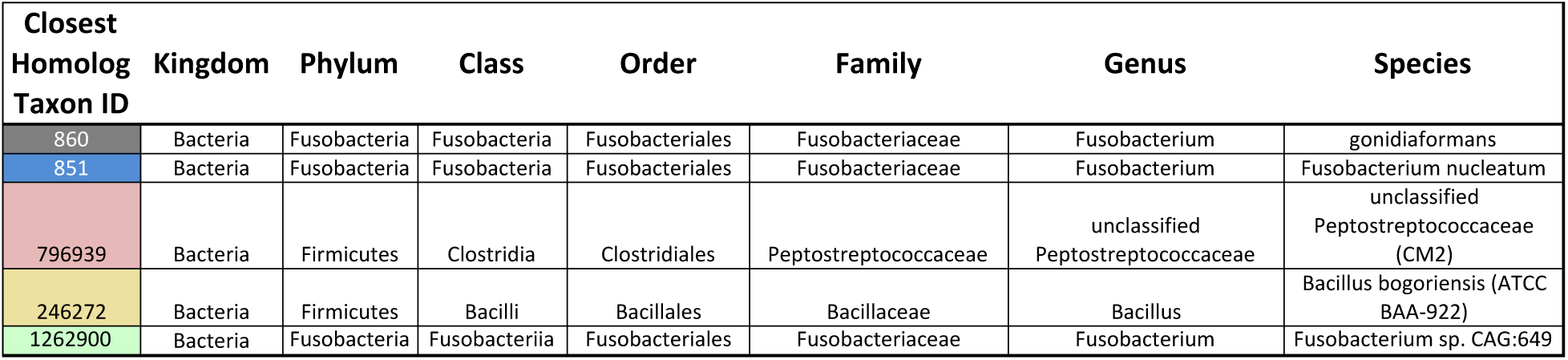
Taxonomic classification of closest homologs to each OTU’s representative sequence (PCA2 phage family).

**SI Figure 3.**
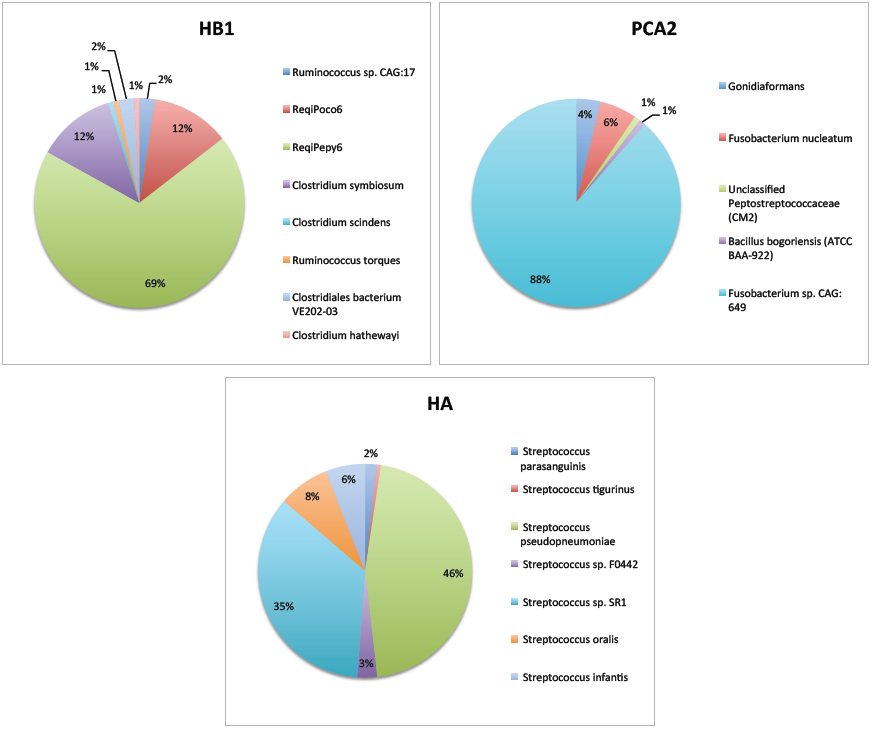
Percentage of HB1, PCA2 and HA phage family OTUs belonging to each taxonomic group identified in SI Figure 2, SI Figure 1, and SI Table 3.

**SI Figure 4.**
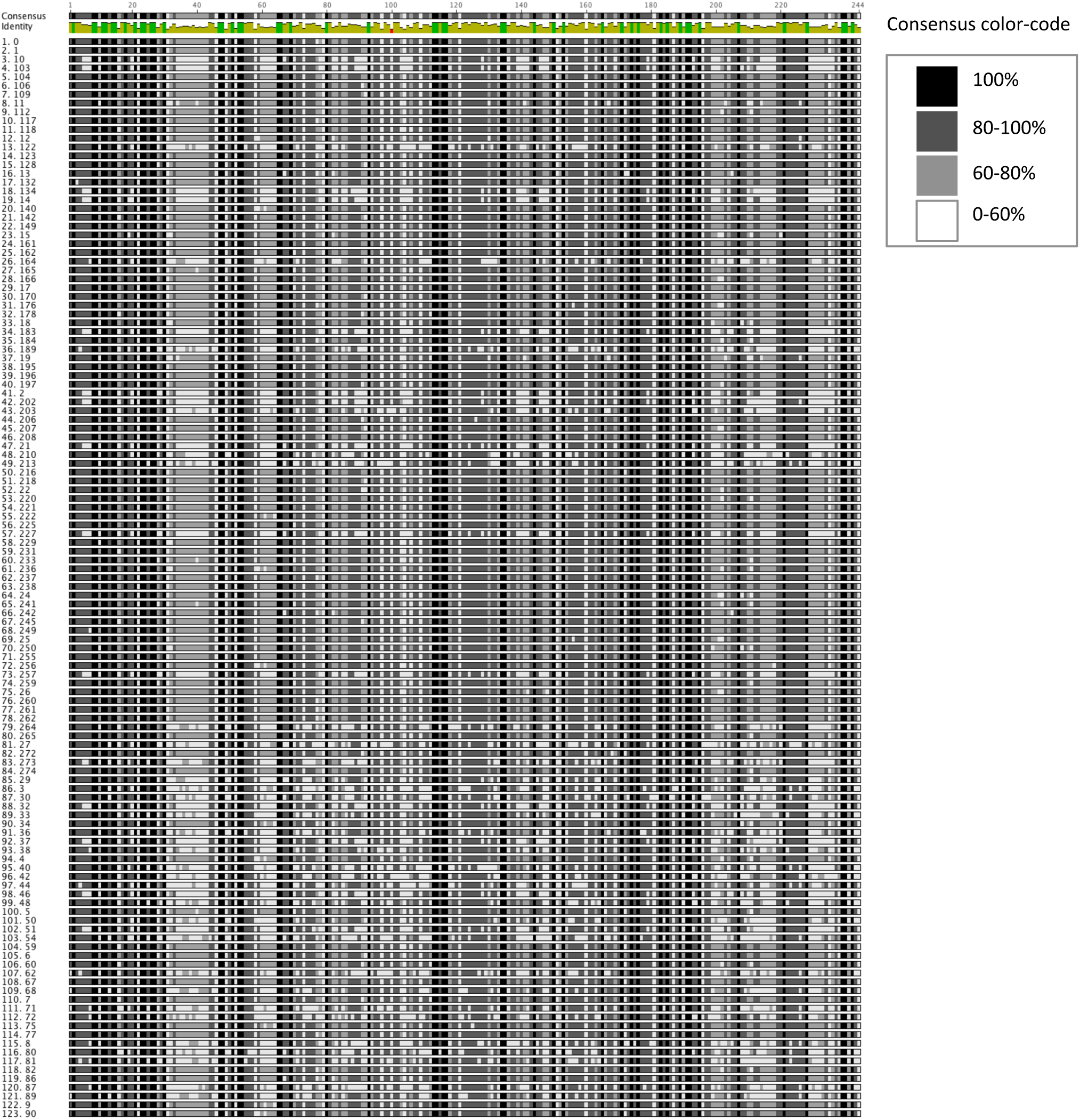
The nucleotide alignment of HB1 phage family OTU representative sequences. Sequences were aligned using Geneious (79). No gaps were introduced. Each base is color-coded according to its relative abundance within a column in the alignment. Conserved bases are black and highly variable sites are denoted in white

## Acknowledgement

We are grateful to members of the Phillips Lab and the Boundaries of Life Initiative for helpful discussions. This study was supported by the National Science Foundation (Graduate Research Fellowship; DGE-1144469), the John Templeton Foundation (Boundaries of Life Initiative; 51250), the National Institute of Health (Maximizing Investigator’s Research Award; RFA-GM-17-002), and the National Institute of Health (Exceptional Unconventional Research Enabling Knowledge Acceleration; R01-GM098465).

